# An in-depth comparison of linear and non-linear joint embedding methods for bulk and single-cell multi-omics

**DOI:** 10.1101/2023.04.10.535672

**Authors:** Stavros Makrodimitris, Bram Pronk, Tamim Abdelaal, Marcel Reinders

## Abstract

Multi-omic analyses contribute to understanding complex biological processes, but also to making reliable predictions about, for example, disease outcomes. Several linear joint dimensionality reduction methods exist, but recently neural networks are more commonly used to embed different-omics into the same non-linear manifold. We compared linear to non-linear joint embedding methods using bulk and single-cell data. For modality imputation, non-linear methods had a clear advantage. Comparisons in downstream supervised tasks lead to the following insights: First, concatenating the principal components of each modality is a competitive baseline for multi-modal prediction. If only one modality was available at test time, joint embeddings yielded significant performance improvements with respect to a unimodal predictor. Second, imputed omics profiles can be fed to classifiers trained on real data with limited performance drops. Overall, the product-of-experts architecture performed well in most tasks while a common encoder of concatenated modalities performed poorly.

---

In the past years there has been a tendency to produce more and more multi-modal-omics data [1], i.e. to measure multiple different data modalities from the same sample. This has enabled the discovery of complex biological mechanisms and lead to deeper understanding of biological processes (e.g. [2, 3]). For example, with joint profiling of genetic and transcriptomic data from the same individuals we can uncover eQTLs (expression quantitative trait loci), i.e. genetic variants that influence the expression of particular genes [4].

A prime example of such multi-modal datasets is The Cancer Genome Atlas (TCGA), which contains mutation, gene expression, DNA methylation, and copy number profiles from the same tumor sample for thousands of primary tumors from different cancer types [5]. More recently, advances in single-cell technology have enabled the simultaneous profiling of different-omics from the same single cell, for example gene expression and DNA methylation [6] or gene expression and protein expression [7]. A review of multi-modal single-cell technologies can be found in [8].

This rapid progress in our ability to generate data has naturally increased the need for computational tools to deal with the challenges of analyzing increasing amounts of multi-modal data. One popular class of such tools is called joint dimensionality reduction or joint embedding and involves projecting all modalities into the same lower-dimensional space [9–11]. The ”joint space” then encodes the information shared by all modalities and filters out modality-specific signals. This not only reduces the effects of experimental noise, but also helps us uncover relationships between modalities.

Several joint embedding methods have been proposed and many of them have been especially designed for-omics data. The majority of such methods are extensions of single modality dimensionality reduction such as factor analysis or principal components analysis. For example, MOFA+ [10] is a generalization of probabilistic PCA for multi-modal data, as it uses variational inference to find a linear projection that minimizes the total reconstruction error of all modalities. Through the use of appropriate prior distributions, MOFA+ also tries to learn a ”sparse” space with a small number of factors and a small number of features contributing to each factor. Other methods do not focus on reconstruction, but to find projections that maximize a criterion. For instance, MCIA [11] maximizes the covariance between the input profiles and the latent representation, while AJIVE [12] finds the directions of maximum variance for the concatenated principal components of all modalities.

Recently, the use of neural networks and deep learning for dimensionality reduction has gained popularity in several domains, including computational biology and mainly single cell transcriptomics [13]. This is motivated by the fact that neural networks can identify non-linear patterns in the data which are expected to be present in-omics data (e.g. synthetic lethality [14] or enhancer synergy [15]). A widely-used model in such settings is the Variational AutoEncoder (VAE) [16], which uses a neural network to learn a probabilistic mapping from an input profile to a latent variable (encoder), while a second network (decoder) learns the inverse mapping (from the latent variables to the input). VAEs have also been used in multi-modal settings, for example to learn a joint embedding of gene and protein expression profiles from CITE-Seq data [17].

Advances in multi-modal deep learning from other fields, such as computer vision can also be applied in computational biology. One such model, also using VAEs, is called Product of Experts (PoE) autoencoder [18]. It uses one encoder for each input modality which are then combined to yield the final joint embedding. The formulation of PoE makes it usable in settings where some modalities are not observed. The Mixture of Experts (MoE) [19] autoencoder modified the way the single-modality embeddings are combined for the joint embedding, leading to improved performance with respect to PoE, although the superiority of MoE is not that well-established in the machine learning literature [20]. Both models have been applied in single-cell multi-omic datasets (e.g. [21, 22]). For example, Minoura et al. [22] used MoE to identify regulatory relationships between gene expression and chromatin accessibility, as well as to predict one modality from the other.

The large number of available methods calls for comparisons between them. Cantini et al. compared nine *linear* joint embedding methods at various tasks using both bulk and single-cell data [9]. The main finding of this study was that there was large variability on the ranking of methods depending on the task, but a few methods, such as MCIA [11] tended to rank near the top for most tasks. Another observation made by the authors was that most of the tested methods were designed for bulk data, but nevertheless also performed reasonably well on single-cell data. Next to being resctricted to linear methods, a limitation of this study is the authors used a single-cell dataset of only 206 cells, which is too small to be representative of modern single-cell experiments, as current single-cell technologies enable multi-omic single-cell profiling at much larger scales. For example, with SNARE-Seq it is possible to measure gene expression and chromatin accessibility in over ten thousand cells [23], and a recent CITE-Seq study profiled gene and protein expression in more than half a million cells in a single experiment [24]. Therefore, possible differences in scalability among these methods were not taken into account. In another study, Brombacher et al. compared the performance of several deep learning-based joint representation learners as a function of the number of cells in the datasets [25]. This study was focused on single-cell data and did not compare to any well-established linear methods.

In this paper, we compare several neural network architectures for joint representation learning to each other as well as to two popular linear methods that also showed promise according to Cantini et al. [9]. We evaluate the models’ ability to impute missing modalities, to learn a coherent latent space for all modalities, and to perform well on downstream tasks using both bulk and single-cell data. Moreover, when appropriate, we employ simple baselines that do not make any use of joint dimensionality reduction to put the observed performances into perspective.

## Results

### Non-linear methods show superior imputation on TCGA

We first compared two linear joint embedding methods, MCIA ([11], Figure 1a) and MOFA+ ([10], Figure 1b) against four non-linear, neural-network-based joint dimensionality reduction methods using the TCGA data. We included two simple non-linear architectures as baseline: concatenated VAE (ccVAE, Figure 1c) works by concatenating the modalities and feeding them to a common encoder. When imputing one modality from the other, all inputs of the missing modality are set to zero as in [17]. Cross-Generating VAE (CGVAE, Figure 1d) uses separate encoders and learns a joint space by a) forcing each encoder’s output to be able to reconstruct all modalities and b) an additional loss term penalizing the difference between the learnt embeddings of each modality based on Wasserstein distance (Online Methods). We also bench-marked two existing VAE-based joint embedding methods, namely product of experts (PoE, [18], Figure 1e) and mixture of experts (MoE, [19], Figure 1f). MCIA and MOFA+ cannot be readily applied to new unseen samples (out-of-sample extension), but since they are linear methods, we enabled this by fitting linear mappings from the input to the embedding space and vice-versa. See Online Methods for more details about all methods and their training.

**Fig. 1.**
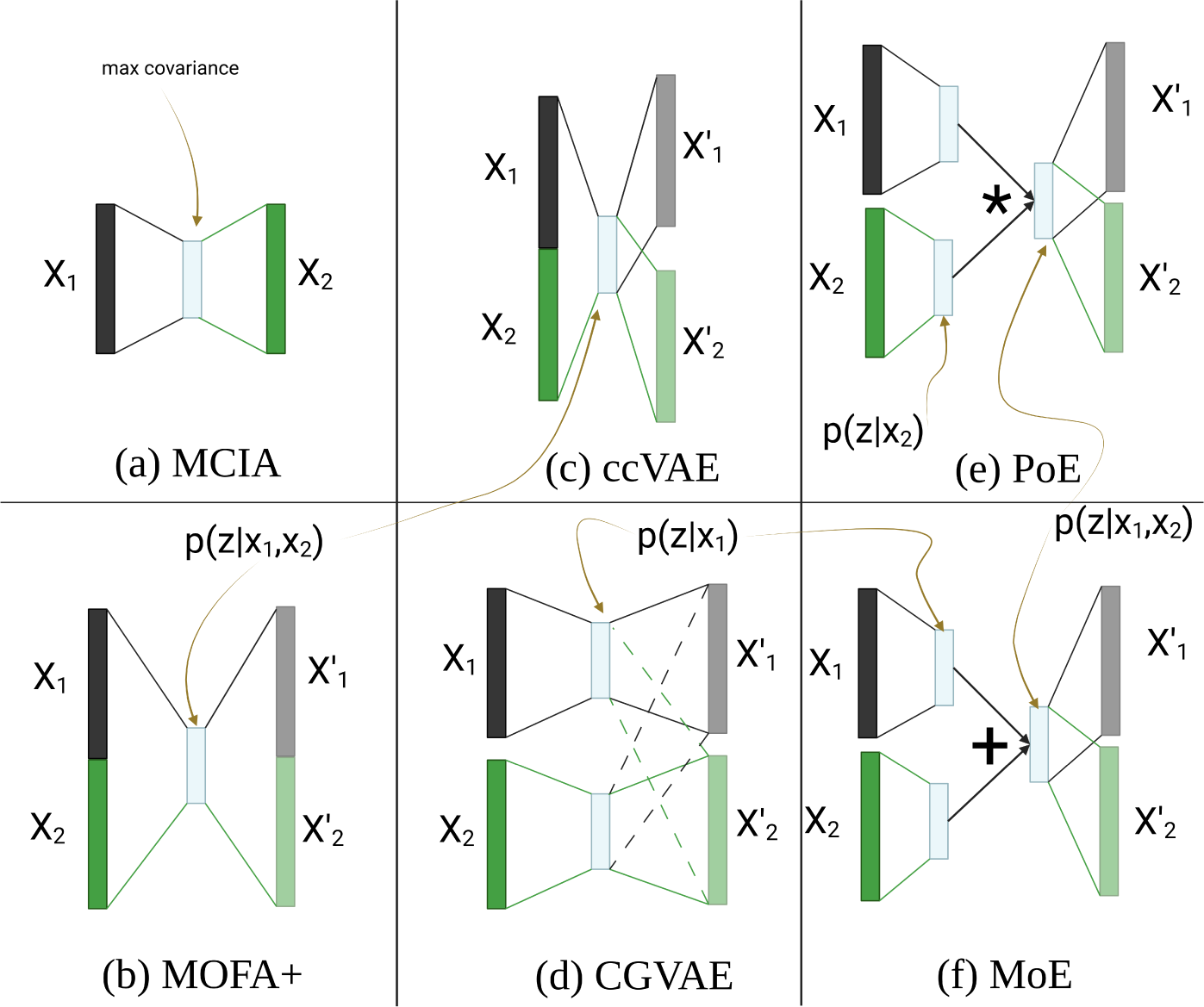
Schematic representation of the six joint representation learning models bench-marked in this paper. On the left, the linear methods: MCIA (a) and MOFA+ (b); in the middle the baseline non-linear methods: ccVAE (c) and CGVAE (d); and on the right the existing non-linear methods: PoE (e) and MoE (f).

Using 6,752 samples from 33 tumor types from the TCGA with gene expression (GE), methylation (ME), and Copy Number Variation (CNV) data available, we learned joint embeddings for GE + ME and GE + CNV, and evaluated the ability to impute a missing modality using a held-out test set of 844 samples (Figure 2a). We also compared against a baseline Generalized Linear Model (GLM) trained to perform regression from one modality to the other without any joint embedding. The results are summarized in Figures 3a and 3b and Tables S1 and S2, where each model’s performance is measured as the log-likelihood of the test data given the model’s predictions (Equation 1, Online Methods).

**Fig. 2.**
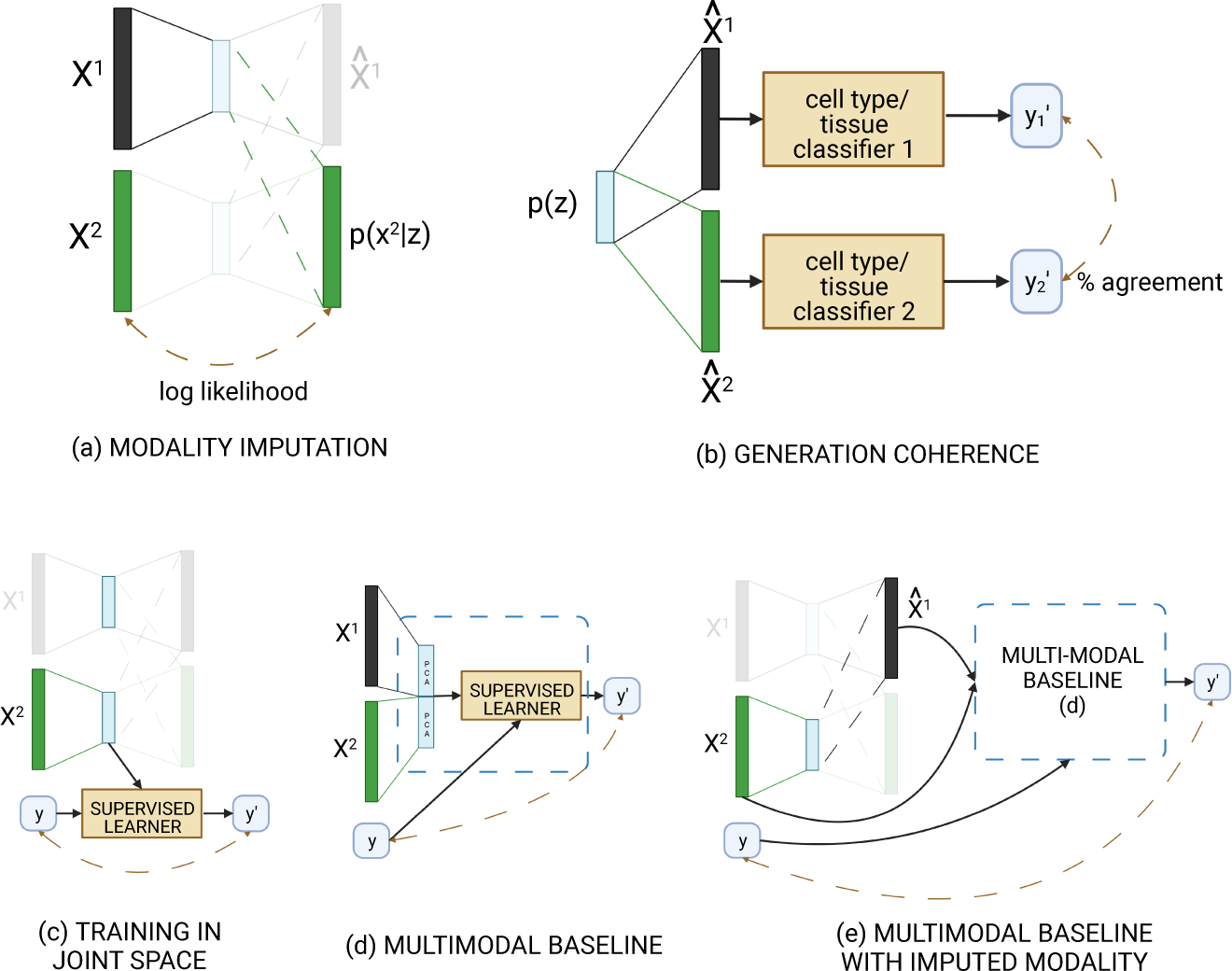
Sketch of our evaluation schemes for (a) modality imputation, (b) generation coherence, (c-e) downstream supervised tasks. Panel (c) shows the training of a classifier in the joint space of one modality, and (d) the baseline method. In (e) we impute the missing modality and feed the measured and the imputed profile into the baseline of panel (d).

**Fig. 3.**
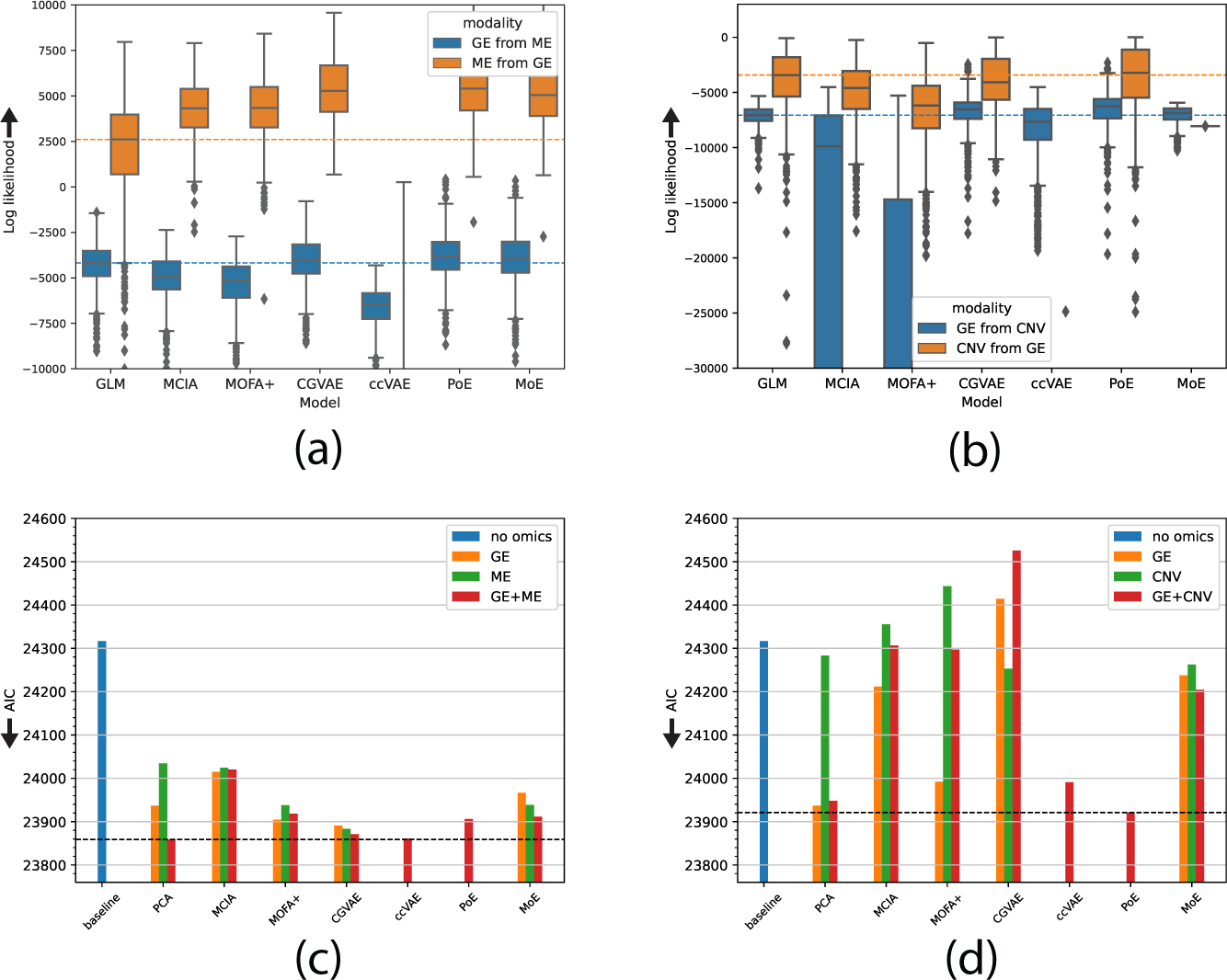
Missing modality imputation and survival analysis performance on the TCGA dataset. (a) Imputing gene expression (GE) from methylation (ME) and vice-versa and (b) gene expression (GE) from copy number variation (CNV) and vice-versa. Performance is measured in both cases as the log-likelihood (*y−*axis, equation 1) of the 844 test samples given the predictions of each model (*x−*axis) for those data (higher is better, signified by upwards-pointing arrows). The dashed horizontal lines represent the performance of the GLM baseline. The median log-likelihood for ccVAE at predicting ME from GE is −70,987.14 and CNV from GE −74,111.33 (not shown to ease visualization of the remaining models). Test samples whose log-likelihood is further than 1.5 times the interquartile range from the median sample are marked as outliers. (c) Predicting progression-free survival using the joint embeddings on the TCGA dataset with GE+ME: The *y−*axis shows the Akaike Information Criterion (AIC, lower is better as signified by downwards arrow) achieved by different models trained in the joint space of GE and ME using either one (GE - orange, ME - green) or two modalities (red). The performance of baseline models using PCA as well as only the covariates (blue) is also shown. (d) As in (c), but for the GE+CNV dataset.

Despite the fact that we also tested hyperparameter configurations without any hidden layers (using a validation set of 844 samples, independent of the training and test sets), we found that, for all four models, the optimal configuration based on the validation set was non-linear with either one (PoE) or two (CGVAE, ccVAE, MoE) hidden layers. The selection of the remaining hyperparameters was highly dependent on the choice of model and dataset (Table S3).

Figures 3a and 3b show that both MCIA and MOFA+ failed to outperform the simple regression baseline in three out of the four cases and only performed well at imputing DNA methylation patterns from gene expression. Furthermore, concatenating a measured modality with a vector of zeros’s for the missing modality and passing the data through ccVAE for imputation lead to very bad performances, considerably worse than all other methods (Figure 3a-b).

CGVAE, PoE and MoE performed better than the other three methods, but only PoE was significantly better than the GLM baseline in all four cases (FWER *<* 0.05, Wilcoxon test). CGVAE and MoE offered signigicant improvements with respect to the baseline in general, but they both failed at predicting CNVs from GE (Figure 3b).

Overall, we observed large improvements in imputation performance with respect to the baseline for the prediction of methylation from gene expression (almost 2-fold increase in log-likelihood), while in most of the remaining cases the differences were marginal (albeit statistically significant). Taken together, we can conclude that PoE is the best performing method in the imputation task.

We additionally compared the coherence of the latent space of the different joint embedding methods by testing whether decodings of the same point in the latent space are classified as the same cancer type by a neural network trained to predict the cancer type using the measured training data (Figure 2b). The predictive performance of the CNV-based cancer type classifier was quite bad with respect to those of the GE- and ME-based classifiers, therefore we restricted the comparison only to the GE and ME dataset (Table S4). Remarkably, PoE and MoE (the top-performing methods at imputation) had the two worst latent space coherence performances (Table 1). The latent space of ccVAE was more coherent, giving decodings from the same cancer type in 75% of the cases, second to CGVAE with 81%. MOFA+ ranked third in terms of coherence, performing slightly better than PoE (Table 1). Note that MCIA is not a generative model, so we cannot sample from its latent space and therefore it is not included in this experiment.

**Table 1.** Generation coherence evaluation. Coherence is measured as the fraction of times the decodings of random points from the latent space are from the same class (cancer type for TCGA, cell type for PBMC). Higher is better. Note that MCIA is not generative and cannot be evaluated in this setting.

| model | TCGA (GE+ME) | PBMC |
| --- | --- | --- |
| MCIA | N/A | N/A |
| MOFA+ | 0.53 | 0.70 |
| CGVAE | 0.81 | 0.76 |
| ccVAE | 0.75 | 0.78 |
| PoE | 0.50 | 0.32 |
| MoE | 0.27 | 0.72 |

From these results, we conclude that there is a discrepancy between imputation and coherence as no architecture is very good at both at the same time. On the other hand, we found that in all tasks the top-performing method was a non-linear one.

### Limited practical utility of joint embedding methods for survival analysis

We then tested whether the pretraining of joint dimensionality reduction would lead to performance gains in the supervised downstream task of survival analysis (Figure 2c+d). Specifically, we used the latent features learned by each model to fit a pan-cancer Cox proportional hazards model to predict progression-free survival using age, gender and cancer type as covariates (Online Methods). We tested these methods with either one modality as input (and then take the latent representation related to that modality), as well as with two modalities as input.

As baseline, we used a model with only the covariates mentioned above. Additionally, we compared to three more simple models: two single-omic models based on the 32 principal components of each data modality (PCA, orange and green bars in Figures 3c-d) and a multi-omic model whose input was the concatenation of the 32 principal components of the two modalities (Figure 2d, red PCA bars in Figures 3c-d).

We evaluated using the Akaike Information Criterion (AIC). The AIC measures the quality of a model’s fit to the data by taking the number of model parameters into account, as it is in principle easier for models with more parameters to overfit to the training data. The results of this experiment (the lower the AIC the better) are shown in Figure 3c-d and Tables S5-S6.

Our baseline model using only the patients’ sex, age, and cancer type (and no-omics) information already provided a statistically significant fit (FWER *<* 0.05) compared to a null model with only an intercept (result not shown). The inclusion of the 32 principal components of gene expression or methylation (PCA) lead to large improvements (Figure 3c). In fact, combining the principal components of these modalities was the best performance we could achieve. PCA on the copy number data only yielded a minor reduction in AIC with respect to the baseline (Figure 3d). Gene expression is the most predictive single modality of the three in this task.

Cox models trained on the latent space of joint embedding methods using gene expression and methylation (GE+ME) did improve upon the baseline, but were all outperformed by PCA, with ccVAE, CGVAE, and PoE ranking second, third, and fourth respectively when both modalities are included (Figure 3c). MCIA had the worst performance in this setting, but still provided a significant fit (FWER *<* 0.05).

As for models trained on the latent space of gene expression and copy number (GE+CNV, Figure 3d), joint pre-training sometimes proved detrimental as CGVAE provided a very poor fit, comparable to what could be expected by chance (FWER *>* 0.05) and MOFA+ and MCIA barely outperformed the baseline. However, PoE did outperform the baseline methods and had the best performance in this dataset, with the baseline PCA second and ccVAE ranking third (Figure 3d).

When we restrict our comparison to methods that only use one modality, we hypothesized that a joint embedding will be beneficial, as the joint pretraining can provide additional information from other modalities. For gene expression (the most informative single modality), this was not in general the case, as only two of the four joint embedding methods (MOFA+ and CGVAE) trained on the joint space of gene expression and methylation (GE+ME) outperformed training only on gene expression data (GE). Cox models that use the gene expression data embedded in the GE+CNV joint space performed worse than the Cox model trained on the PCA of the GE data. This implies that the common information between gene expression and copy number does not relate to disease progression, possibly because CNV data have relatively small prognostic power (Figure 3d).

On the other hand, training a model on the joint methylation space (GE+ME) does improve the AIC performance when comparing to training on only methylation data (ME) for all joint models (Figure 3c). Moreover, gene expression informed by methylation outperformed gene expression informed by copy number for all four models (Extended Figure 1). These results indicate that the joint embedding space of gene expression and copy number does encode information about the metastatic potential of tumors.

### Neural architectures impute and scale better on a large single-cell dataset

We then compared the same six joint embedding methods on a CITE-Seq dataset profiling Peripheral Blood Mononuclear Cells (PBMCs) from 8 different individuals [26]. We used 6 individuals for training (126,424 cells), one for validation (17,205 cells), and one for testing (18,135 cells). The dataset also contains cell type annotations for each cell. These annotations are provided at three different levels of granularity: Level 1 is the most coarse labeling with 8 different classes, Level 2 uses 31 classes, and Level 3 is the most fine-grained labeling with 58 classes.

The best configurations for the non-linear models were as follows: MoE architecture was composed of three layers and the selected CGVAE and ccVAE models had two layers. None of the selected models used dropout or batch normalization. Surprisingly, the PoE configuration with the lowest validation loss was linear, i.e. had no hidden layers and included dropout (Table S7). We tried to train MCIA on this training set with 750GB of RAM but it did not terminate due to insufficient memory after several computation hours so it is not included in this comparison.

We first compared the methods on missing modality imputation, using a similar GLM-based baseline method as for the TCGA data (Figure 2a). All methods, including the baseline, performed similarly at imputing RNA (RNA) from protein expression (ADT), with MoE having a slight edge over the rest, followed by PoE (Figure 4a, Table S8). Pairwise Wilcoxon rank sum tests showed that the performance differences were statistically significant (FWER *<* 0.05) despite the small absolute differences in median. By inspection of the imputation performance per (level 2) cell type and clustering the cell types based on their median imputation log-likelihood, we found two distinct clusters of cell types (Extended Figure 2). That means that there is one group of cell types for which all models can perform good imputations. These include erythrocytes, naive T cells, B cells and natural killer cells. On the other hand, for the other cluster, which includes monocytes, proliferating T cells, and hematopoietic progenitors, imputing RNA from ADT is harder. Nevertheless, MoE has a clear edge on the cells of that cluster with respect to all other methods (Extended Figure 2).

**Fig. 4.**
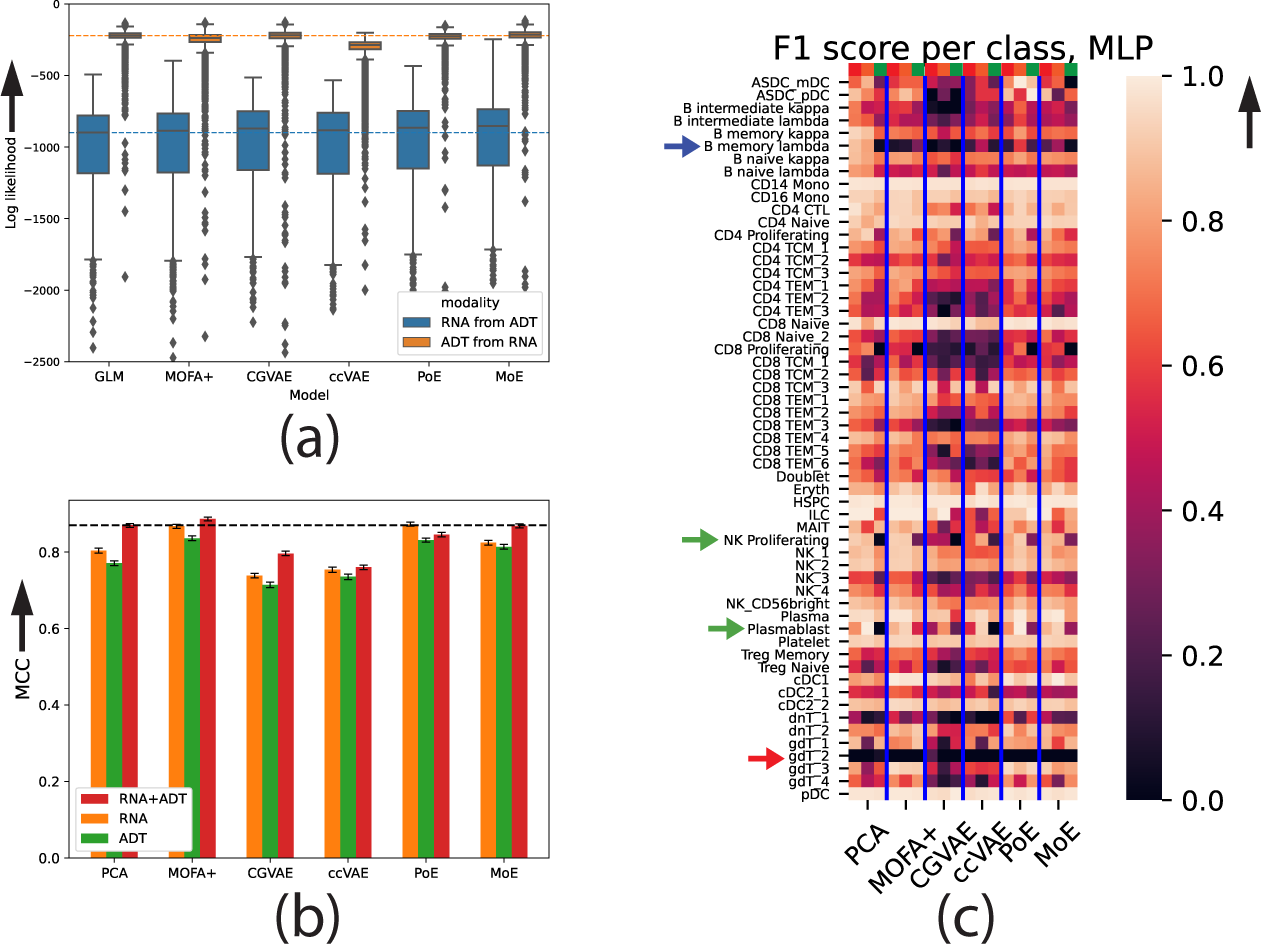
Evaluation on the CITE-Seq dataset. (a) Missing modality imputation performance for gene expression (RNA) from protein expression (ADT) and vice-versa. Performance is measured as the log-likelihood (*y−*axis, equation 1) of the test samples (cells) given the predictions of each model (*x−*axis) for those data (higher is better). The distribution of the per-cell log-likelihoods is shown. The dashed horizontal lines represent the performance of the baseline GLM. Cells further than 1.5 times the interquartile range from the median are marked as outliers. (b) Cell type classification performance (MCC, *y−*axis, higher is better) achieved by training a multilayer perceptron (MLP) in the joint space of the different models when using: only gene expression (RNA, orange), only protein expression (ADT, green), and both RNA and ADT data (red). The error bars denote 95% confidence intervals calculated by bootstrapping the test cells 100 times. (c) Per-class (cell type) performance of the same classifiers as in (b). Brighter colors denote a higher per-class F1 score and therefore better performance. For each model we show three columns (RNA+ADT-red, RNA only-orange, and ADT only-green, signified by top row). Arrows show the cell types highlighted in the results. Note that class CD4+ Tem 4 is not present in the test data and therefore not shown in the per-class evaluations (because its precision and recall is always 0 and the F1 score is thus undefined), but it was taken into account when calculating the MCC in (b).

Imputation of ADT from RNA followed the pattern of the TCGA experiments, with MOFA+ and ccVAE performing worse than the GLM baseline, and the other non-linear methods having a slight edge. In this experiment, we observed much smaller variance in the imputation log-likelihood compared to the RNA (Figure 4a and Extended Figure 3), with most of the outlier cells (i.e. cells for which the imputation is less accurate than the rest) being erythrocytes. To evaluate generation coherence (Figure 2b), we built neural networks that predict the level 2 cell type from the RNA or the ADT data (Table S4). We found that CGVAE and ccVAE, had the highest performance (as for the TCGA dataset). MoE and MOFA+ had improved coherence with respect to the TCGA dataset, but were still inferior. PoE again performed badly in this task (Table 1).

### Multi-modal pre-training partly compensates for an unmeasured modality at test time

We also compared the pre-trained models on the supervised task of cell type classification (Figure 2c), using the level 3 labels containing in total 58 different classes (cell types). Using the same data split as for the imputation, we used the training set of 6 individuals to train both the joint embedding methods and two cell type classifiers: a linear SVM and a two-layer MLP (Online Methods). We evaluated on the cells of a left-out individual. As in the TCGA survival analysis experiment, we also used a baseline method that projects each modality to its 32 principal components and trained the same classifiers (Figure 2d).

We again found that joint unsupervised pre-training on the same dataset does not yield any considerable advantage to downstream performance if both modalities are available at test time, as concatenating the PCs of gene and protein expression (RNA+ADT) gave competitive performance using both classifiers (Figure 4b, Extended Figure 4, Table S9). PoE and MoE embeddings did perform marginally better than this baseline using the SVM classifier and MOFA+ and MoE performed marginally better or equally using the MLP (Table S9). CGVAE and ccVAE did not perform well in this task. We also observed that the MLP (Figure 4b-c) outperformed the SVM classifier (Extended Figure 4) regardless of the input data, which shows that at least some of the classes are not linearly separable and require non-linear modeling. We thus focus on the MLP classifier onwards.

Then we evaluated all methods when only one modality is available at test time and compared the models that are pre-trained in the joint space against a classifier trained on the PCs of the single modality. In this setting, there was a drop in the performances with respect to the case when both modalities are always available. However, we observed considerable improvements caused by the joint pre-trainining using MOFA+, PoE, and MoE, as demonstrated in Figure 4b (orange and green bars). A MLP trained on the RNA latent space of PoE or MOFA+ performed only slightly worse than the best performance that was achieved when having two modalities available. When measuring only protein expression, we obtained similar results, with joint pre-training using MOFA+, PoE, and MoE providing a significant performance gain with respect to the baseline (Figure 4b). Furthermore, ADT data embedded on the joint RNA+ADT space gave worse performance than RNA data embedded on this space for all methods. This implies that the two joint spaces are not equivalent and that RNA is ’the dominant modality’, although we corrected for the fact that RNA has more features than ADT (Online Methods).

To gain a better understanding of the performance differences between the different models, we examined the performance per class as measured by the F1 score (Figure 4c). First, we observed that a very rare class of *γδ* T cells (gdT cell 2) is very hard to differentiate among other T cell types. Only the PCA of protein expression data and CGVAE with both modalities was able to do better than random on this class (red arrow in Figure 4c). *λ* memory B cells are predicted accurately using PCA on the RNA data, but the performance of the ADT principal components and all joint embedding methods is poor (blue arrow in Figure 4c). This implies that the information needed to distinguish this cell type is only in the RNA and therefore it is not present in the joint space (RNA+ADT), so joint pre-training is detrimental for the performance in this specific class. On the other hand, proliferating natural killer cells and plasmablasts (green arrows) are also only predicted well by RNA, but embedding RNA on the joint RNA+ADT space of PoE and MoE did not hurt the performance for those classes.

### Imputed modalities are useful for cell type classification

Next, we considered whether it is possible to use the joint embedding methods to impute the missing modality and then classify the test cells using the MLP on the concatenated principal components of the two modalities (one measured, one imputed, Figure 2e). When only RNA is available for the test data, we used MOFA+, PoE, and MoE to impute the corresponding ADT profiles, and combined the measured RNA and imputed ADT profiles to be used by the cell type MLP classifier operating on the PCs of both modalities. Figure 5a shows that for all three methods this approach led to higher MCC than a simple classifier trained on only RNA data. However, this approach did not outperform classification models trained on the joint space of RNA (Figure 2c) that also only require RNA measured at test time (denoted as joint unimodal in Figure 5).

**Fig. 5.**
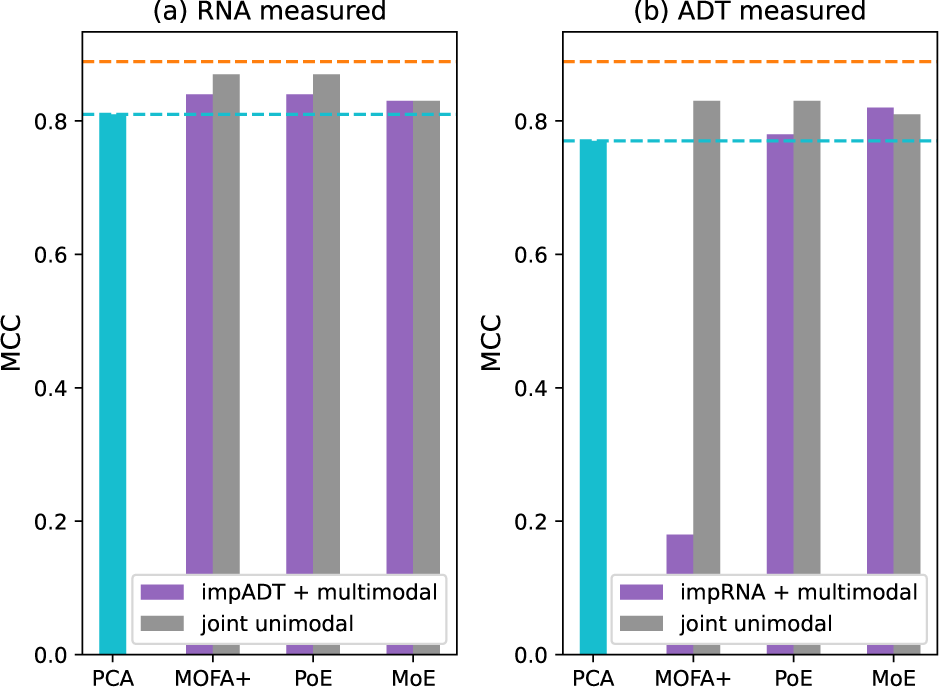
Comparison of training a cell type classifier in joint space (joint unimodal, Figure 2c) versus using the joint space to impute a missing modality and using a classifier trained on both modalities (impRNA/ADT + multimodal, Figure 2e). Panel (a) shows the case when only RNA is available at test time and (b) when only ADT measurements are available. Performance is quantified by the Matthews Correlation Coefficient (MCC, higher is better). The light blue bar and dashed line correspond to the performance of a MLP trained only on the measured modality, while the dashed orange line shows the highest performance achieved by any model that used both measured modalities (0.89).

If we only use ADT data on the test set, we find that imputing RNA and then using a multi-modal model was detrimental for MOFA+, but doing so using PoE and MoE did lead to higher classification performance with respect to a model trained only on ADT (Figure 5b). Only for MoE, which performed best at the imputation task, a multi-modal model with imputed RNA was even slightly better than training a model on the ADT joint space.

Because we actually have both modalities measured for all test cells, we can compare the predictions made by the multi-modal MLP when using either both measured profiles or when one of them is imputed (Table S10). When we use imputed protein profiles alongside gene expression, the multi-modal classifier predicts the same class as when using the measured protein profiles for 89.1%, 89.6% and 88.7% of the cells when the imputation is done using MOFA+, PoE, and MoE, respectively. The corresponding agreement rates when we combine measured protein profiles with imputed RNA profiles are 11.5%, 83.6% and 86.2%, respectively (Table S10).

We additionally used this set-up to further test how realistic the generated profiles are. If we have measured RNA at test time, we can impute the corresponding protein profiles, project them to their 32 PCs and feed them into the ADT classifier (Table S10). Although this doesn’t have any practical utility and leads to worse performances than using the multi-modal classifier, we found that the predictions of the ADT classifier after performing this imputation matched the predictions of the same classifier made using the measured data. As shown in Table S10, the imputed and measured profiles gave the same prediction in 77.3%, 74.7% and 73.3% of the test cells for MOFA+, PoE and MoE, respectively. When doing the reverse experiment, i.e. imputing RNA from protein profiles, projecting to the RNA PCs and feeding into the RNA classifier, the agreement rates were 6.2%, 71.3% and 77.5% for MOFA+, PoE and MoE, respectively. These agreement rates are slightly smaller than those we found when feeding one measured and one imputed modality into the multi-modal, but they are still significantly higher than what we would expect if the classifier was randomly guessing when predicting using imputed profiles. The relatively large correspondence between predictions made with measured and imputed profiles is additional evidence for the high imputation quality of non-linear joint embedding methods. On the other hand, ADT profiles imputed using MOFA+ had high agreement measured profiles, but RNA imputation was much less successful in this task.

## Discussion

We presented a comparison of established linear joint embedding methods to novel neural-network-based methods that can learn non-linear mappings in both bulk and single-cell multi-omics data. We also included simple appropriate baselines that do not employ any joint dimensionality reduction in each experiment, which are often omitted in similar studies (e.g. [9]).

We found that non-linear methods developed in other fields (PoE and MoE) generally outperformed the linear and simple non-linear ones at imputing missing modalities. On the other hand, these methods underperformed with respect to baseline non-linear methods in terms of generation coherence. Regarding downstream supervised tasks, we observed that joint embedding can lead to improved performance when only a single modality is available in the test data, which verifies previous results for linear methods [27]. If data from both modalities are available at test time, joint embedding did not provide a significant advantage in the tasks that we tested here.

Interestingly, we additionally showed that joint embedding methods can be used to impute missing modalities which can be fed to a multi-modal classifier trained only on real data. This means that the tested models are able to generate realistic enough-omic profiles. On the other hand, these profiles were projected to the principal components space before being fed to the classifier, which provides an additional denoising step. In most cases, this approach of imputing and classifying with two modalities was worse than training a model in the joint space using one modality. This is not surprising, as imputation errors are bound to be propagated into the classifier despite the denoising. On the other hand this approach did show improvement over using a supervised model trained on a single modality.

To make sure we fairly compare all methods, we performed an extensive hyperparameter search to find the best settings for each one. We used a held-out validation set to calculate the validation loss for each hyperparameter combination and select the optimal combination. The performances of the methods were estimated using another held-out test set comprising previously unseen data points. If, for some reason, the validation loss is not predictive of the performance at a specific downstream task - as we have previously shown can be the case for VAEs trained on RNA-Seq data [28] - then the chosen hyperparameters for a given model might not be the optimal ones for the downstream task. Our results here hint that this might be the case in our setting too. Many complex models did well at imputing one modality from the other, which is part of the loss function for training them, but showed less impressive results in other downstream tasks. Therefore, if the goal of learning a joint embedding is performing well at a specific downstream task, we recommend to employ a semi-supervised training scheme, where labeled data are used during the embedding learning process. With these labeled data one can simultaneously minimize the sum of the joint embedding loss and the supervised loss of a model trained on the target task with inputs from the joint latent space as in [29, 30].

Examining the results on generation coherence, we found that CGVAE and ccVAE did better than PoE and MoE on both bulk and single-cell data, while they typically underperformed in the other tasks. ccVAE uses a single encoder for the concatenation of both modalities, which might be beneficial for generation coherence, as the latent space is directly and concurrently influenced by matched samples from all modalities. However, the architecture of CGVAE is identical to that of MoE and PoE with separate encoders per modality. What makes CGVAE different from these models is an additional loss term that penalizes the Wasserstein distance between the posterior distribution of the latent variables given a pair of input modalities. This encourages the latent embeddings from both modalities to be the same for the same input sample. Coherence of MoE and PoE could be potentially be improved by adding such a loss term, but that also introduces an additional hyperparameter to weigh the contribution of that loss component with respect to the total loss. Here, we used the CGVAE model as a simple baseline, thus we did not tune this hyperparameter value.

Our results showed that our adaptation of MOFA+ [10] with out-of-sample extension was in some cases competitive with state-of-the-art neural-based embedding methods, especially in the CITE-Seq data. The addition of the linear regression on the latent variables can be seen as an additional regularization step [31] which stabilizes the model and might partly explain the good performance. In addition, MOFA+ has the advantage that it provides useful diagnostic messages about the input data as well as the learnt space. For instance, it automatically removes latent factors that explain too little variance.

Cantini et al. concluded that many of the linear joint embedding methods they tested performed well on single-cell data [9]. Here, we used a more recent single-cell dataset with many more cells and found that one of the best methods in the benchmark of Cantini et al., MCIA, was not even possible to train despite using a considerable amount of computational power. This points to an advantage of VAEs: they have been designed to work with stochastic or batch gradient descent to accommodate large training datasets [16]. Of the nine methods tested by Cantini et al. [9], only MOFA+ [10] and scikit-fusion [32] offer batch training mode and GPU acceleration and are therefore applicable to the latest generation of single-cell datasets with at least tens of thousands of cells. Many of the other methods, such as MCIA, include an eigendecomposition or singular value decomposition step, which can get very expensive for large sample and feature sizes, in terms of both time and memory. Accelerated versions of these operations (e.g. [33]) might alleviate this burden.

An important consequence of our experimental set-up is, that the number of data points used to train the joint embedding methods is equal to the number of points used to train the downstream supervised models. In practice, additional unlabeled multi-modal data might be available, which can be used during the learning of the joint space. In such cases, especially as the amount of unlabeled data increases, we expect an additional benefit of joint embedding, even if both modalities are available at test time.

As a final note, it is worth pointing out that next to learning a joint embedding space, it is interesting to learn a latent representation of the signal that is unique to each modality. In the linear setting, this has been achieved by AJIVE [12], which treats directions with significant variance that are orthogonal to the joint space as the modality-specific or ”individual” space. When dealing with non-linear embeddings, however, finding these individual spaces is more complicated. One possible solution could be to use adversarial losses to ”force” part of the latent space to not be useful for reconstructing the other modality.

## Online Methods

### Notation

We consider datasets of *N* samples where we measure *M* different modalities (gene expression, DNA methylation etc.) for each sample *i* with the *m*-th modality represented by a feature vector 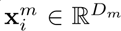. We seek to find a joint latent representation for the 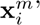 denoted by **z***_i_ ∈* R*^d^*, where *d < D_m_* for all *m*, which contains the common information of all modalities. The *j*-th element of 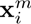, is denoted as 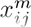. The joint embeddings are learned using a model - for example an encoder neural network - that learns the parameters *θ_i_* of the distribution 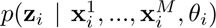. For instance, the most common choice for the posterior of *z* is a Gaussian distribution with diagonal covariance, in which case *θ_i_* corresponds to *d* mean and *d* standard deviation values for each sample that are learned by the encoder. The entire set of samples from the *m*-th modality is represented by the matrix 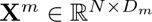.

In the case of autoencoder-like methods, the learned latent representations are used during training to reconstruct the input data by learning the parameters *ϕ* of the ’generative’ distribution 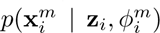 for each modality *m*. The reconstruction quality of each modality from the latent space is assessed using the log-likelihood (*LL*) as defined in equation 1. As it is common in the literature, we assume that the likelihood factorizes over the features. The reconstructed modality is denoted as 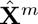 consisting of row vectors 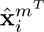 In the following, we omit the conditioning on *θ* and *ϕ* for simplicity.

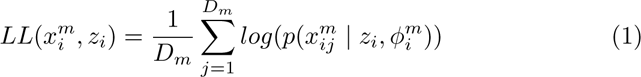

### Algorithms for joint representation learning

We compared two linear methods, MOFA+ [10] and MCIA [11], two existing neural architectures (product of experts [18] and mixture of experts [19]) and two simpler, baseline non-linear joint representation learners. The six methods, visualized in Figure 1, are briefly described below. Details about training and hyperparameter optimization are provided in the Supplementary Methods.

### Linear methods

MOFA+ infers a common low-dimensional latent space from a set of high-dimensional data modalities [10]. Each modality is mapped into the common latent space using a projection matrix **W***^m^*, such that 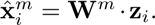 MOFA+ makes use of prior distributions for **W** that ensure both shrinkage and differential activity of each latent dimension across modalities and employs variational inference to find the matrices that minimize the sum of the per-modality negative log-likelihoods [10]. MOFA+ only support Gaussian, Poisson, and Bernoulli likelihoods.

MCIA attempts to find projections that maximize the covariance between features of each modality and a common latent space called the reference structure [11].

To allow for out-of-sample generalization for these methods, we need the projection (encoding) matrices that map a sample 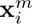to **z***_i_* and, as we are interested in imputing one modality from the other, we also need the inverses of these linear mappings (decoding matrices). Neither of these methods provide all the necessary matrices in their standard implementation, so we estimated them as follows: Using the training data, we fit a linear regression to predict **z***_i_* from 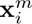, for each modality *m* and for the concatenation of all modalities. The weights of those fitted regressions from the input data to the MOFA+/MCIA output are used as encoding matrices to embed unseen samples. We obtain the decoding matrices similarly (predicting 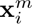 from **z***_i_*), but this time we use multivariate Generalized Linear Models (GLMs) to deal with the different distribution of each data modality (see Datasets and preprocessing).

### Baseline non-linear embedding methods

The simplest method to combine different modalities is to concatenate the features into one vector 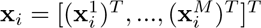 and use them to train a VAE (*ccVAE*). The concatenated vector **x***_i_* is fed into a probabilistic encoder that learns *q*(**z***_i_ |* **x***_i_*) and separate decoders are used to decode each modality from **z***_i_*. The model is trained by minimizing the sum of the negative log likelihood plus a Kullback-Leibler divergence term (*KL*) between *q*(**z***_i_ |* **x***_i_*) and the prior distribution of the latent variables *p*(**z**) (equation 2). We used Gaussian distributions with diagonal covariance for both the prior and the posterior of **z** [16] Note that totalVI [17] uses this architecture, but models the data differently.

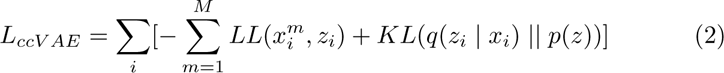

Another simple non-linear joint representation learning architecture consists of one VAE for each data source. To force the encoders to embed the data in a common space we want the decoders to be able to reconstruct any modality from the latent representation of any encoder. To do so, we minimize the sum of the reconstruction losses for all possible combinations of input and output (reconstructed) modalities. We further add a loss term penalizing the second-order Wasserstein distance between the posterior distribution of **z** given each input data source. This encourages the output of different encoders (i.e. the learned embeddings of the same sample) to be similar. The loss function of this model, which we call Cross-Generating Variational Autoencoder (CGVAE), is shown in Equation 3, where *W*_2_ is the second-order Wasserstein distance.

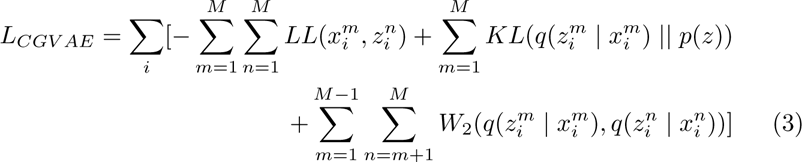

### Existing non-linear embedding methods

The Product of Experts approach (PoE) [18] also uses a single VAE per data modality, but combines all the per-modality latent representations into one final posterior distribution for **z**. This distribution 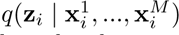 is given by multiplying the individual densities 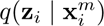 with each other as well as with the prior *p*(**z**). If *q* and *p* are chosen to be Gaussian, then the resulting product is also a Gaussian (up to a normalizing constant). In practice, PoE models are trained by sampling latent vectors from both the joint 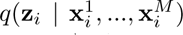, as well as the individual 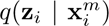, passing each of those through all decoders and summing the different reconstruction losses. For more details on the loss function of PoE, see the original publication [18].

Mixture of Experts (MoE) [19] uses the mixture of the individual densities so that 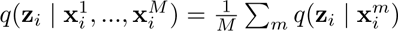. In addition, following the original publication [19], when training MoE, we replaced the Gaussian priors and posteriors of *z* with Laplace distributions and employed the DReG gradient estimator [34] to reduce the variance of the gradient estimates.

### Training details Hyperparameters

We trained all neural network models using PyTorch and the Adam optimizer with batch size of 64. We used the common choice of *N* (0*, I*) for *p*(**z**) and a normal distribution for the variational posterior *q*(**z** *|* **x**) for CGVAE, ccVAE and PoE. For MoE, we followed the original publication and used Laplace distributions for both the prior and the posterior. We employed a grid search for each model and in each of the three datasets to find the optimal combination of the following hyperparameters:

- learning rate (1*e −* 3 or 1*e −* 4)
- dimensionality of *z* (32 or 64)
- encoder hidden layers (none, 128, 256, 256-256, 256-128)
- dropout probability (10% or no dropout)
- use of batch normalization (yes or no)

In each case the decoder architecture was symmetric to that of the encoder. MoE has one more hyperparameter: the number of samples (K) drawn from the posterior of **z**. For that we tried K=10, or 20. During the grid search, we trained each configuration for a maximum of 500 epochs and applied early stopping if the validation loss did not improve for more than 0.5% for longer than 10 epochs.

### Implementation details

When we pass an omic profile from modality *m* 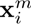 into the encoder network, we obtain the latent representation of that sample in the joint space of modality *m* via the distribution 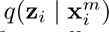. One exception to that is ccVAE, which uses concatenated profiles from all modalities. Therefore, for ccVAE, to calculate the parameters of 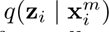, we set the second modality to a vector of zero’s.

When both modalities are present, obtaining 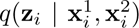 is straightforward for ccVAE (profiles are concatenated and passed through the encoder). For PoE, it is obtained by multiplying the densities 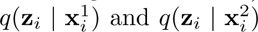, which gives us a new Gaussian distribution. We can then obtain a single embedding for sample *i* by taking the mean of this distribution.

In the case of MoE, however, this is not straightforward. 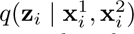 is a mixture of 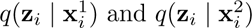 and the mean of that mixture distribution might be a vector that is very improbable by both single-modality posteriors. During training, this is ammended by drawing multiple samples from the variational posterior, but for our downstream analyses we need a single feature vector per sample. CGVAE suffers from a similar issue, as its formulation does not provide a method to obtain a single joint vector based on both modalities. For these two models, we obtained a latent representation based on both modalities by concatenating the mean vectors of 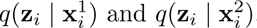.

### MCIA

We used the R package omicade4 to run MCIA. Training this joint embedding on the PBMC data was not possible due to extreme memory requirements (*>* 750GB), but we ran into convergence problems in the TCGA datasets too. To keep the number of latent factors comparable with the neural networks, we ran MCIA with 32 and 64 latent factors, but both of these setting lead to runtime convergence errors and did not yield any output in both GE+ME and GE+CNV. We then started decreased the number of factors from 32 in steps of 4 until we could obtain an output.

### MOFA+

We ran MOFA+ with 32 and 64 latent factors in CPU mode (i.e. without the GPU acceleration feature), using the MOFA2 R package. All the settings were left to their default values, except for the early stopping parameter (called ”convergence mode”), which we set to ’medium’. MOFA+ includes a post-processing step where factors not explaining any variance, which is why we got models with fewer than 64 factors in Tables S3 and S7. We applied the built-in ”select model” function to select the best of the two models.

### Datasets and pre-processing

#### TCGA dataset

The TCGA dataset [35] consists of 8,440 samples of tumors - including a few adjacent normal samples as well for which gene expression (GE), copy number variation (CNV) and DNA methylation (ME) are measured for 33 different tumor types. For GE, we used the batch-corrected data [36], standardized to zero mean and unit variance per gene with a Gaussian likelihood. For CNV, we used the per-gene copy number data [37] estimated using GISTIC2 [38]. The copy number estimates were discretized using the GISTIC2 thresholds into one of five possible categories: homozygous deletion, heterozygous deletion, normal copy number, low-level amplification, and high-level amplification. As the data are discrete, we used the categorical log-likelihood. For ME [39], we restricted to samples measured with Illumina 450K chip. We grouped CpG sites that lie within 1,000 bp from the transcription start site of the 24,994 protein-coding genes (Ensembl version 79), averaging the beta values within each group ignoring missing values. If all CpG beta values of a group were missing for a particular sample, we set that feature value to 0. Finally, to accomodate the use of a beta log-likelihood, we replaced all zero’s with a small number (*ɛ* = 10*^−^*^6^) and all one’s with 1 *−ɛ*. For each data modality, we selected the 5,000 most variable features based on median absolute deviation. Clinical (meta-)data for the samples were collected from [40].

#### CITE-Seq data

We used a CITE-Seq dataset [26] containing single-cell expression profiles for RNA and surface proteins of peripheral blood cells measured using RNA-Seq and anti-body-derived tags (ADTs) respectively. We log-transformed the RNA count data using a pseudo-count of 1 and modelled them using a negative binomial log-likelihood to account for zero inflation. We used the median absolute deviation to select the 5,000 most variable gene expression features. We retained all 224 protein features measured, then standardized to zero mean and unit variance and used the Gaussian log-likelihood.

### Experiments and evaluation

#### Missing modality imputation

We split each dataset into a training, a validation and a test set. We evaluated the ability of joint embedding methods on predicting (imputing) one data modality from the other, by holding out one modality at a time from the test data (Figure 2a).

We used the log-likelihood of the held-out data given the model predictions as evaluation measure. For example, when imputing, say ME from GE, we give each GE profile (**x***^GE^*) as input to the GE encoder which calculates *q*(**z***_i_ |***x***^GE^*). We take the mean of that distribution as the final embedding vector of sample *i* and feed it to the ME decoder. The log-likelihood of the decoded ME profile then defines the performance. ccVAE requires both modalities to be fed as input, so, similar to [17], we replaced the missing modality with a vector of zeroes and passed the concatenated vector to the encoder. As a baseline imputation method, we trained a GLM to perform regression from one modality to the other using for each modality the same log-likelihood as described in the previous section.

For the TCGA data, the splits (80-10-10%) were stratified per tumor type to account for the vast imbalances between them. We trained then models using two modalities at a time: GE + ME and GE + CNV. As the inter-individual variations are very large in the PBMC dataset, to avoid information leaks stemming from using cells from the same individual during training and testing, we split the cells based on the eight donors. Specifically, we used all cells from 6 individuals for training, 1 individual for validation, and 1 for testing. Although this split is not stratified per cell type, it provides a more realistic and less biased evaluation.

#### Generation coherence

Given a point in the latent space, we expect all decoders to generate instances of each modality with similar properties [19]. We assessed this by randomly drawing from *p*(**z**) and checking whether the decoded profiles reflected ”similar” samples. As these are generated profiles, we cannot check whether they come from the same individual, so we tested whether they resemble profiles from the same cancer type or cell type.

Specifically, we trained five 2-layer perceptrons, one for each modality (GE, ME and CNV for TCGA, RNA and ADT for PBMC), where the input is a modality profile and the target output is the cancer type or the (level 2) cell type, respectively. Again using ME and GE as an example, for each model, we randomly sampled 2,000 points from the prior distribution of **z** and decoded each point using both decoders to generate 2,000 ME and 2,000 GE matched profiles. We then fed the generated profiles into the corresponding perceptron classifier and compared the predictions of the two classifiers for the generations of the same randomly-sampled point in the latent space. Joint decoding quality, also called latent space coherence, was measured as the agreement rate of the two classifiers, i.e. the fraction of random **z**’s whose decoded profiles are predicted to be from the same cancer type. This experiment is illustrated in Figure 2b.

Here, we are not interested in whether the predicted labels of the random points are correct or not, as long as the two perceptrons make the same prediction for profiles generated from the same value of **z**. However, this percent agreement is only meaningful if the perceptrons are well-trained and can accurately discriminate among cancer/cell types. As we see in Table S4, the cancer type predictor trained on CNV data network was not accurate, thus we did not perform this experiment in the GE + CNV dataset.

#### Survival analysis on the TCGA dataset

We compared the latent features obtained from the different joint embedding methods on their ability to predict disease progression. To this end, we performed survival analysis using Cox regression (Figure 2c) with Progression-Free Survival (PFS) as the outcome variable [41]. We used the entire dataset to fit the models and centered and scaled each latent feature to zero mean and unit variance. To compare the generalization capability of the different models, we used the Akaike Information Criterion (AIC) [42], which is defined as 2*k −* 2*ln*(*L^s^*), where *k* is the number of parameters the Cox model has to estimate, and *L^s^* is the model’s likelihood. The model with the lowest AIC value provides the best trade-off between fitting the data well and not using too many parameters. It has been shown that model selection using AIC is asymptotically equivalent to using cross-validation [43]. Therefore, the model with the lowest AIC is considered to generalize best to new, unseen data.

To distil effects of each individual modality, this evaluation took place using the joint embeddings of either both modalities (*q*(**z** *|* **x**_1_, **x**_2_)) or of only one modality at a time (e.g. *q*(**z** *|* **x**_1_)). See Supplementary Methods for more details. We included the patients’ sex (2 one-hot encoded features), age, and cancer type (33 one-hot encoded binary features) as covariates in all models to account for the effects of these variables. A model with only the covariates (i.e. without any-omics features) was used as a baseline. We also compared to three stronger baselines per dataset, namely the 32 principal components of each modality and the concatenation of these pairs of 32 latent features (Figure 2d). The number of principal components was not tuned and was selected to be similar to the dimensionality of the joint embedding spaces.

#### Cell type classification on the PBMC dataset

Because we observed that predicting the level 2 assignments was a relatively easy task in this PBMC dataset (Table S4), we used the more fine-grained level 3 labeling to make the problem more challenging. The train-validation-test split was identical to the one used for the imputation experiment.

We compared the performance of the different embeddings when both modalities are available for all samples, and when both modalities are available for the training and validation data, but only one modality is measured in the test data. For each joint embedding method, we thus obtained three latent vectors: 1) using only RNA, 2) using only proteins and 3) using both RNA and proteins (Supplementary Methods).

We trained two different classifiers for each feature vector: a linear Support Vector Machine (SVM) and a 2-layer perceptron. For the SVM, we chose the best value for the weight of the L2 regularization from the values [10*^−^*^4^, 10*^−^*^3^, 0.01, 0.1, 0.5, 1.0, 2.0, 5.0, 10.0, 20.0] using the validation data and the Matthews Correlation Coefficient (MCC) as criterion. For the perceptron, we set the number of hidden neurons to 64, the learning rate to 0.0001 and trained for 150 epochs minimizing the cross-entropy loss between the ground truth and the predicted cell type labels. The epoch with the lowest validation loss was chosen for evaluating the network.

We additionally built baseline classifiers that did not use any joint embedding, but were trained on a) the 32 principal components of the RNA data, b) the 32 principal components of the ADT data, and c) the concatenation of a and b. We used the same classifiers with the same settings as above and again did not tune the number of principal components for the task.

Finally, Figure 2e shows a third evaluation scheme for cell type classification. In this setting, we again assume that only one modality is measured at test time, but instead of using the joint space of that modality, we used the trained joint embedding methods to impute the missing modality for each cell. The imputed profiles are then projected to the 32 PC space and fed into the baseline multi-modal classifier (Figure 2d).

We then compare the predictions of the multi-modal classifier when using two measured modalities versus one measured and one imputed modality as well as to a classifier trained on the joint space of the measured modality (Figure 2c).

## Supplementary information

Supplementary Material is available.

## Acknowledgments

The results published here are in part based upon data generated by the TCGA Research Network: https://www.cancer.gov/tcga. We would like to thank Mohammed Charrout, Dr. Soufiane Mourragui and Chirag Raman for the useful discussions about generative models and the Convergence Health and Technology program of Erasmus Medical Center and Delft University of Technology for funding part of this work.

## Data Availability

We used publically available data from The Cancer Genome Atlas (https://portal.gdc.cancer.gov/) and Hao et al. (https://atlas.fredhutch.org/nygc/multimodal-pbmc/).

## Code Availability

The code to reproduce the experiments is available at https://github.com/brmprnk/jointomicscomp.

## Extended Data

**Extended Figure 1.**
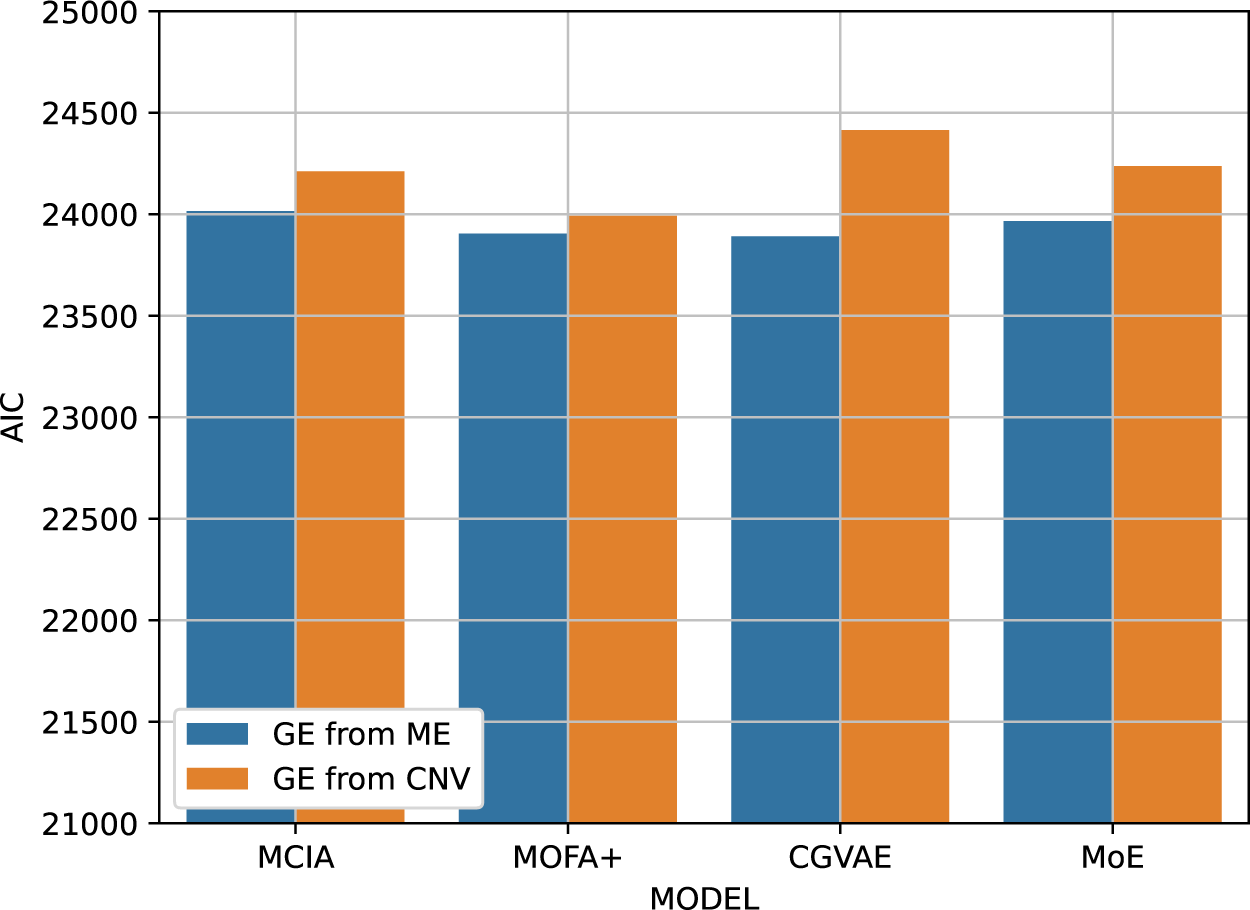
Predictive performance (AIC, *y−*axis) of progression-free survival of gene expression data trained in the joint space of gene expression and methylation (blue) or gene expression and copy number(orange) based on different joint embedding methods (*x−*axis).

**Extended Figure 2.**
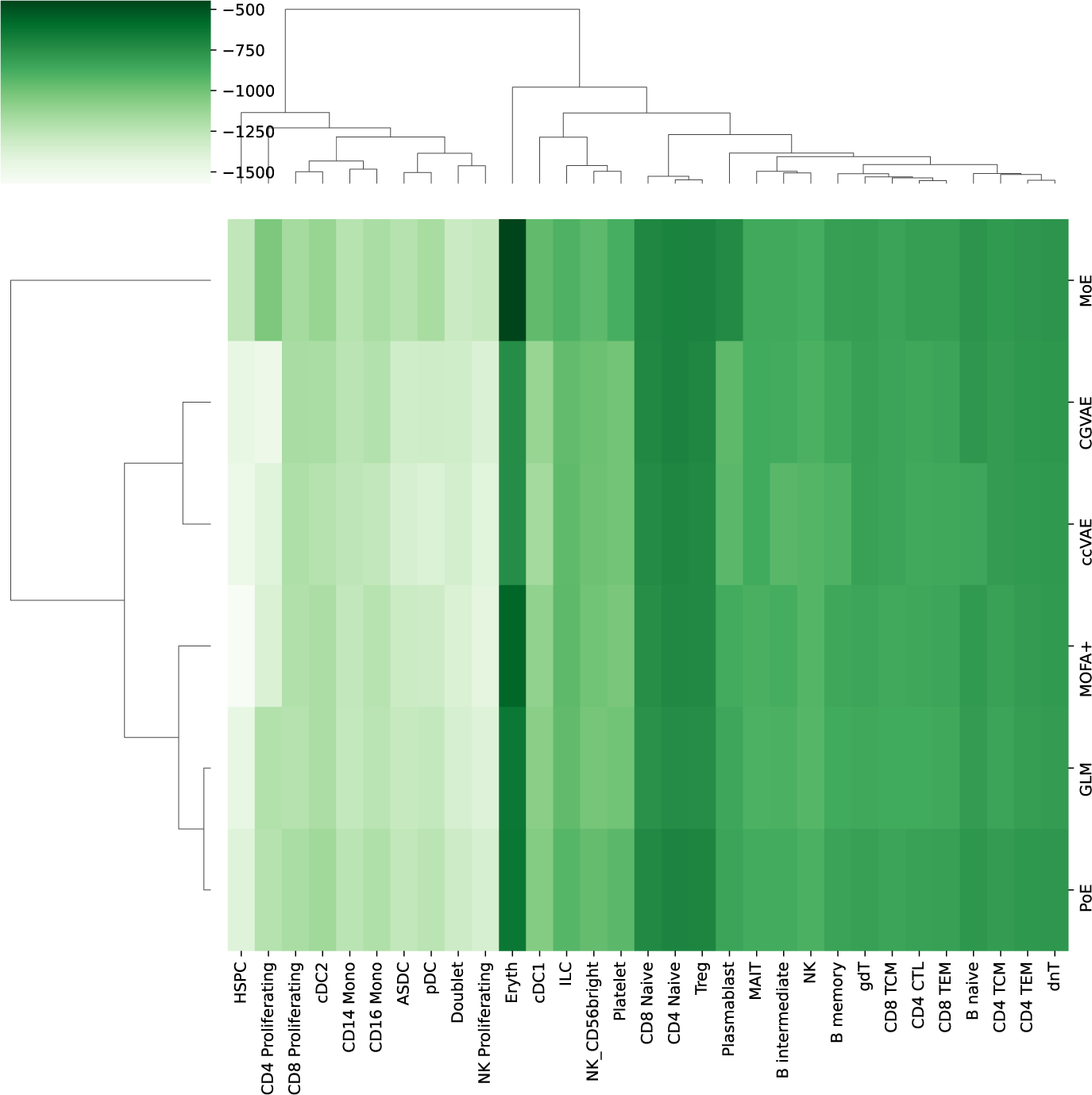
Log likelihood of imputing RNA from ADT for each model. The average log-likelihood is calculated per cancer type for each model. Models and cancer types are clustered so that similar models/cancer types are next to each other. Higher log-likelihood (deeper green) corresponds to better performance.

**Extended Figure 3.**
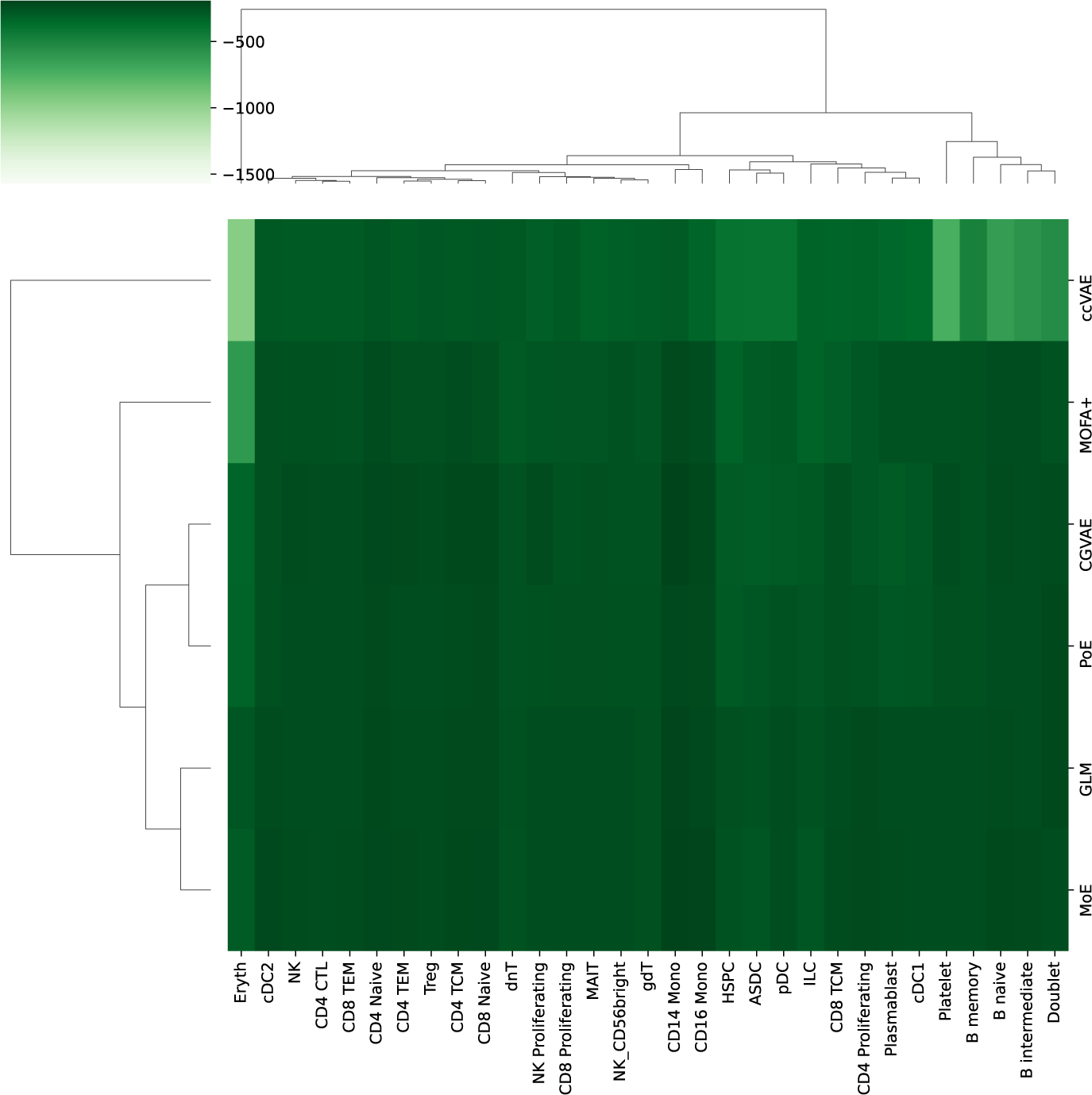
Log likelihood of imputing ADT from RNA for each model. The average log-likelihood is calculated per cancer type for each model. Models and cancer types are clustered so that similar models/cancer types are next to each other. Higher log-likelihood (deeper green) corresponds to better performance.

**Extended Figure 4.**
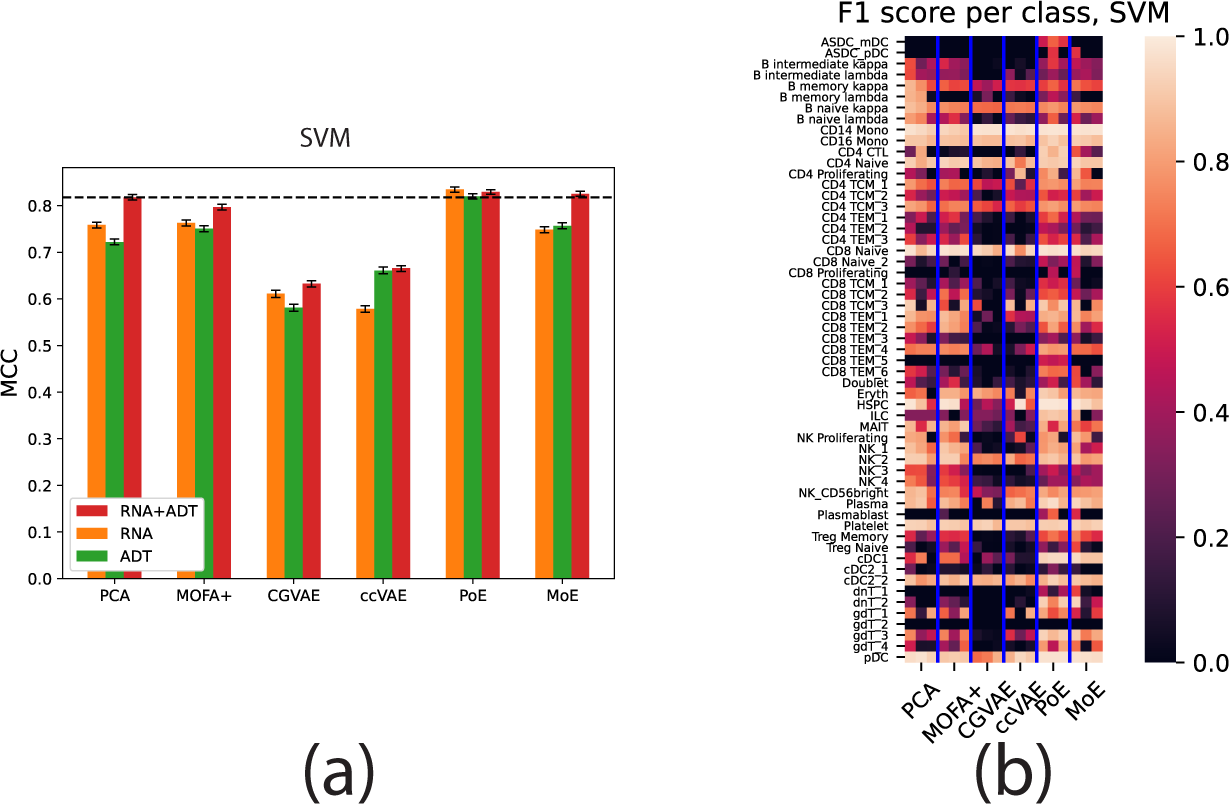
Cell type classification performance when using a linear SVM classifier instead of a MLP. (a) Classification performance (MCC, *y−*axis, higher is better) achieved by training a linear SVM in the joint space of the different models when using: only gene expression (RNA, orange), only protein expression (ADT, green), and both RNA and ADT data (red). The error bars denote 95% confidence intervals calculated by bootstrapping the test cells 100 times. (b) Per-class (cell type) performance of the same classifiers. Brighter colors denote a higher per-class F1 score and therefore better performance. For each model we show three columns (RNA+ADT, RNA only and ADT only). Note that class CD4+ Tem 4 is not present in the test data and therefore not shown in the per-class evaluations (because its precision and recall is always 0 and the F1 score is thus undefined), but it was taken into account when calculating the MCC in (a).

## References

[1] Krassowski, M., Das, V., Sahu, S.K., Misra, B.B.: State of the Field in Multi-Omics Research: From Computational Needs to Data Mining and Sharing. Front Genet 11, 610798 (2020)

[2] Li, J., Shi, M., Ma, Z., Zhao, S., Euskirchen, G., Ziskin, J., Urban, A., Hallmayer, J., Snyder, M.: Integrated systems analysis reveals a molecular network underlying autism spectrum disorders. Mol Syst Biol 10(12), 774 (2014)

[3] Frattini, V., Trifonov, V., Chan, J.M., Castano, A., Lia, M., Abate, F., Keir, S.T., Ji, A.X., Zoppoli, P., Niola, F., Danussi, C., Dolgalev, I., Porrati, P., Pellegatta, S., Heguy, A., Gupta, G., Pisapia, D.J., Canoll, P., Bruce, J.N., McLendon, R.E., Yan, H., Aldape, K., Finocchiaro, G., Mikkelsen, T.e G.G., Bigner, D.D., Lasorella, A., Rabadan, R., Iavarone, A.: The integrated landscape of driver genomic alterations in glioblastoma. Nat Genet 45(10), 1141–1149 (2013)

[4] Neavin, D., Nguyen, Q., Daniszewski, M.S., Liang, H.H., Chiu, H.S., Wee, Y.K., Senabouth, A., Lukowski, S.W., Crombie, D.E., Lidgerwood, G.E., ndez, D., Vickers, J.C., Cook, A.L., Palpant, N.J., bay, A., Hewitt, A.W., Powell, J.E.: Single cell eQTL analysis identifies cell type-specific genetic control of gene expression in fibroblasts and reprogrammed induced pluripotent stem cells. Genome Biol 22(1), 76 (2021)

[5] Weinstein, J.N., Collisson, E.A., Mills, G.B., Shaw, K.R., Ozenberger, B.A., Ellrott, K., Shmulevich, I., Sander, C., Stuart, J.M., Chang, K., Creighton, C.J., Davis, C., Donehower, L., Drummond, J., Wheeler, D., Ally, A., Balasundaram, M., Birol, I., Butterfield, S.N., Chu, A., Chuah, E., Chun, H.J., Dhalla, N., Guin, R., Hirst, M., Hirst, C., Holt, R.A., Jones, S.J., Lee, D., Li, H.I., Marra, M.A., Mayo, M., Moore, R.A., Mungall, A.J., Robertson, A.G., Schein, J.E., Sipahimalani, P., Tam, A., Thiessen, N., Varhol, R.J., Beroukhim, R., Bhatt, A.S., Brooks, A.N., Cherniack, A.D., Freeman, S.S., Gabriel, S.B., Helman, E., Jung, J., Meyerson, M., Ojesina, A.I., Pedamallu, C.S., Saksena, G., Schumacher, S.E., Tabak, B., Zack, T., Lander, E.S., Bristow, C.A., Hadjipanayis, A., Haseley, P., Kucherlapati, R., Lee, S., Lee, E., Luquette, L.J., Mahadeshwar, H.S., Pantazi, A., Parfenov, M., Park, P.J., Protopopov, A., Ren, X., Santoso, N., Seidman, J., Seth, S., Song, X., Tang, J., Xi, R., Xu, A.W., Yang, L., Zeng, D., Auman, J.T., Balu, S., Buda, E., Fan, C., Hoadley, K.A., Jones, C.D., Meng, S., Mieczkowski, P.A., Parker, J.S., Perou, C.M., Roach, J., Shi, Y., Silva, G.O., Tan, D., Veluvolu, U., Waring, S., Wilkerson, M.D., Wu, J., Zhao, W., Bodenheimer, T., Hayes, D.N., Hoyle, A.P., Jeffreys, S.R., Mose, L.E., Simons, J.V., Soloway, M.G., Baylin, S.B., Berman, B.P., Bootwalla, M.S., Danilova, L., Herman, J.G., Hinoue, T., Laird, P.W., Rhie, S.K., Shen, H., Triche, T., Weisenberger, D.J., Carter, S.L., Cibulskis, K., Chin, L., Zhang, J., Getz, G., Sougnez, C., Wang, M., Saksena, G., Carter, S.L., Cibulskis, K., Chin, L., Zhang, J., Getz, G., Dinh, H., Doddapaneni, H.V., Gibbs, R., Gunaratne, P., Han, Y., Kalra, D., Kovar, C., Lewis, L., Morgan, M., Morton, D., Muzny, D., Reid, J., Xi, L., Cho, J., DiCara, D., Frazer, S., Gehlenborg, N., Heiman, D.I., Kim, J., Lawrence, M.S., Lin, P., Liu, Y., Noble, M.S., Stojanov, P., Voet, D., Zhang, H., Zou, L., Stewart, C., Bernard, B., Bressler, R., Eakin, A., Iype, L., Knijnenburg, T., Kramer, R., Kreisberg, R., Leinonen, K., Lin, J., Liu, Y., Miller, M., Reynolds, S.M., Rovira, H., Shmulevich, I., Thorsson, V., Yang, D., Zhang, W., Amin, S., Wu, C.J., Wu, C.C., Akbani, R., Aldape, K., Baggerly, K.A., Broom, B., Casasent, T.D., Cleland, J., Creighton, C., Dodda, D., Edgerton, M., Han, L., Herbrich, S.M., Ju, Z., Kim, H., Lerner, S., Li, J., Liang, H., Liu, W., Lorenzi, P.L., Lu, Y., Melott, J., Mills, G.B., Nguyen, L., Su, X., Verhaak, R., Wang, W., Weinstein, J.N., Wong, A., Yang, Y., Yao, J., Yao, R., Yoshihara, K., Yuan, Y., Yung, A.K., Zhang, N., Zheng, S., Ryan, M., Kane, D.W., Aksoy, B.A., Ciriello, G., Dresdner, G., Gao, J., Gross, B., Jacobsen, A., Kahles, A., Ladanyi, M., Lee, W., Lehmann, K.V., Miller, M.L., Ramirez, R., tsch, G., Reva, B., Sander, C., Schultz, N., Senbabaoglu, Y., Shen, R., Sinha, R., Sumer, S.O., Sun, Y., Taylor, B.S., Weinhold, N., Fei, S., Spellman, P., Benz, C., Carlin, D., Cline, M., Craft, B., Ellrott, K., Goldman, M., Haussler, D., Ma, S., Ng, S., Paull, E., Radenbaugh, A., Salama, S., Sokolov, A., Stuart, J.M., Swatloski, T., Uzunangelov, V., Waltman, P., Yau, C., Zhu, J., Hamilton, S.R., Getz, G., Sougnez, C., Abbott, S., Abbott, R., Dees, N.D., Delehaunty, K., Ding, L., Dooling, D.J., Eldred, J.M., Fronick, C.C., Fulton, R., Fulton, L.L., Kalicki-Veizer, J., Kanchi, K.L., Kandoth, C., Koboldt, D.C., Larson, D.E., Ley, T.J., Lin, L., Lu, C., Magrini, V.J., Mardis, E.R., McLellan, M.D., McMichael, J.F., Miller, C.A., O’Laughlin, M., Pohl, C., Schmidt, H., Smith, S.M., Walker, J., Wallis, J.W., Wendl, M.C., Wilson, R.K., Wylie, T., Zhang, Q., Burton, R., Jensen, M.A., Kahn, A., Pihl, T., Pot, D., Wan, Y., Levine, D.A., Black, A.D., Bowen, J., Frick, J., Gastier-Foster, J.M., Harper, H.A., Helsel, C., Leraas, K.M., Lichtenberg, T.M., McAllister, C., Ramirez, N.C., Sharpe, S., Wise, L., Zmuda, E., Chanock, S.J., Davidsen, T., Demchok, J.A., Eley, G., Felau, I., Ozenberger, B.A., Sheth, M., Sofia, H., Staudt, L., Tarnuzzer, R., Wang, Z., Yang, L., Zhang, J., Omberg, L., Margolin, A., Raphael, B.J., Vandin, F., Wu, H.T., Leiserson, M.D., Benz, S.C., Vaske, C.J., Noushmehr, H., Knijnenburg, T., Wolf, D., Van’t Veer, L., Collisson, E.A., Anastassiou, D., Ou Yang, T.H., Lopez-Bigas, N., Gonzalez-Perez, A., Tamborero, D., Xia, Z., Li, W., Cho, D.Y., Przytycka, T., Hamilton, M., McGuire, S., Nelander, S., Johansson, P., rnsten, R., Kling, T., Sanchez, J.: The Cancer Genome Atlas Pan-Cancer analysis project. Nat Genet 45(10), 1113–1120 (2013)

[6] Angermueller, C., Clark, S.J., Lee, H.J., Macaulay, I.C., Teng, M.J., Hu, T.X., Krueger, F., Smallwood, S., Ponting, C.P., Voet, T., Kelsey, G., Stegle, O., Reik, W.: Parallel single-cell sequencing links transcriptional and epigenetic heterogeneity. Nat Methods 13(3), 229–232 (2016)

[7] Stoeckius, M., Hafemeister, C., Stephenson, W., Houck-Loomis, B., Chattopadhyay, P.K., Swerdlow, H., Satija, R., Smibert, P.: Simultaneous epitope and transcriptome measurement in single cells. Nat Methods 14(9), 865–868 (2017)

[8] Dimitriu, M.A., Lazar-Contes, I., Roszkowski, M., Mansuy, I.M.: Single-Cell Multiomics Techniques: From Conception to Applications. Front Cell Dev Biol 10, 854317 (2022)

[9] Cantini, L., Zakeri, P., Hernandez, C., Naldi, A., Thieffry, D., Remy, E., Baudot, A.: Benchmarking joint multi-omics dimensionality reduction approaches for the study of cancer. Nat Commun 12(1), 124 (2021)

[10] Argelaguet, R., Arnol, D., Bredikhin, D., Deloro, Y., Velten, B., Marioni, J.C., Stegle, O.: MOFA+: a statistical framework for comprehensive integration of multi-modal single-cell data. Genome Biology 21(111) (2020). https://doi.org/10.1186/s13059-020-02015-1

[11] Meng, C., Kuster, B., Culhane, A.C., Gholami, A.M.: A multivariate approach to the integration of multi-omics datasets. BMC Bioinformatics 15, 162 (2014)

[12] Feng, Q., Jiang, M., Hannig, J., Marron, J.S.: Angle-based joint and individual variation explained. Journal of Multivariate Analysis 166, 241–265 (2018). https://doi.org/10.1016/j.jmva.2018.03.008

[13] Lopez, R., Regier, J., Cole, M.B., Jordan, M.I., Yosef, N.: Deep generative modeling for single-cell transcriptomics. Nat Methods 15(12), 1053–1058 (2018)

[14] O’Neil, N.J., Bailey, M.L., Hieter, P.: Synthetic lethality and cancer. Nat Rev Genet 18(10), 613–623 (2017)

[15] Choi, J., Lysakovskaia, K., Stik, G., Demel, C., Soding, J., Tian, T.V., Graf, T., Cramer, P.: Evidence for additive and synergistic action of mammalian enhancers during cell fate determination. eLife 10, 65381 (2021). https://doi.org/10.7554/eLife.65381

[16] Kingma, D.P., Welling, M.: Auto-Encoding Variational Bayes. 2nd International Conference on Learning Representations (2013) arXiv:1312.6114 [stat.ML]

[17] Gayoso, A., Steier, Z., Lopez, R., Regier, J., Nazor, K.L., Streets, A., Yosef, N.: Joint probabilistic modeling of single-cell multi-omic data with totalVI. Nat Methods 18(3), 272–282 (2021)

[18] Wu, M., Goodman, N.D.: Multimodal generative models for scalable weakly-supervised learning. CoRR abs/1802.05335 (2018) arXiv:1802.05335

[19] Shi, Y., Siddharth, N., Paige, B., Torr, P.H.S.: Variational Mixture-of-Experts Autoencoders for Multi-Modal Deep Generative Models (2019)

[20] [20] Kutuzova, S., Krause, O., McCloskey, D., Nielsen, M., Igel, C.: Multimodal Variational Autoencoders for Semi-Supervised Learning: In Defense of Product-of-Experts (2021). https://openreview.net/forum?id=aHfiIow3m

[21] [21] Inecik, K., Uhlmann, A., Lotfollahi, M., Theis, F.: Multicpa: Multimodal compositional perturbation autoencoder. bioRxiv (2022) https://www.biorxiv.org/content/early/2022/07/10/2022.07.08.499049.full.pdf. https://doi.org/10.1101/2022.07.08.499049

[22] [22] Minoura, K., Abe, K., Nam, H., Nishikawa, H., Shimamura, T.: Scmm: Mixture-of-experts multimodal deep generative model for single-cell multiomics data analysis. bioRxiv (2021) https://www.biorxiv.org/content/early/2021/02/19/2021.02.18.431907.full.pdf. https://doi.org/10.1101/2021.02.18.431907

[23] Chen, S., Lake, B.B., Zhang, K.: High-throughput sequencing of the transcriptome and chromatin accessibility in the same cell. Nat Biotechnol 37(12), 1452–1457 (2019)

[24] Stephenson, E., Reynolds, G., Botting, R.A., Calero-Nieto, F.J., Morgan, M.D., Tuong, Z.K., Bach, K., Sungnak, W., Worlock, K.B., Yoshida, M., Kumasaka, N., Kania, K., Engelbert, J., Olabi, B., Spegarova, J.S., Wilson, N.K., Mende, N., Jardine, L., Gardner, L.C.S., Goh, I., Horsfall, D., McGrath, J., Webb, S., Mather, M.W., Lindeboom, R.G.H., Dann, E., Huang, N., Polanski, K., Prigmore, E., Gothe, F., Scott, J., Payne, R.P., Baker, K.F., Hanrath, A.T., Schim van der Loeff, I.C.D., Barr, A.S., Sanchez-Gonzalez, A., Bergamaschi, L., Mescia, F., Barnes, J.L., Kilich, E., de Wilton, A., Saigal, A., Saleh, A., Janes, S.M., Smith, C.M., Gopee, N., Wilson, C., Coupland, P., Coxhead, J.M., Kiselev, V.Y., van Dongen, S., Bacardit, J., King, H.W., Rostron, A.J., Simpson, A.J., Hambleton, S., Laurenti, E., Lyons, P.A., Meyer, K.B.c M.Z., Duncan, C.J.A., Smith, K.G.C., Teichmann, S.A., Clatworthy, M.R., Marioni, J.C., ttgens, B., Haniffa, M., Baker, S., Bradley, J.R., Dougan, G., Goodfellow, I.G., Gupta, R.K., Hess, C., Kingston, N., Lehner, P.J., Matheson, N.J., Owehand, W.H., Saunders, C., Smith, K.G.C., Summers, C., Thaventhiran, J.E.D., Toshner, M., Weekes, M.P., Bucke, A., Calder, J., Canna, L., Domingo, J., Elmer, A., Fuller, S., Harris, J., Hewitt, S., Kennet, J., Jose, S., Kourampa, J., Meadows, A., O’Brien, C., Price, J., Publico, C., Rastall, R., Ribeiro, C., Rowlands, J., Ruffolo, V., Tordesillas, H., Bullman, B., Dunmore, B.J., Fawke, S., f, S., Hodgson, J., Huang, C., Hunter, K., Jones, E., Legchenko, E., Matara, C., Martin, J., O’Donnell, C., Pointon, L., Pond, N., Shih, J., Sutcliffe, R., Tilly, T., Treacy, C., Tong, Z., Wood, J., Wylot, M., Betancourt, A., Bower, G., De Sa, A., Epping, M., Huhn, O., Jackson, S., Jarvis, I., Marsden, J., Nice, F., Okecha, G., Omarjee, O., Perera, M., Richoz, N., Sharma, R., Turner, L., De Bie, E.M.D.D., Bunclark, K., Josipovic, M., Mackay, M., Michael, A., Rossi, S., Selvan, M., Spencer, S., Yong, C., Ansaripour, A., Mwaura, L., Patterson, C., Polwarth, G., Polgarova, P., Stefano, G.D., Allison, J., Butcher, H., Caputo, D., Clapham-Riley, D., Dewhurst, E., Furlong, A., Graves, B., Gray, J., Ivers, T., Kasanicki, M., Gresley, E.L., Linger, R., Meloy, S., Muldoon, F., Ovington, N., Papadia, S., Phelan, I., Stark, H., Stirrups, K.E., Townsend, P., Walker, N., Webster, J.: Single-cell multi-omics analysis of the immune response in COVID-19. Nat Med 27(5), 904–916 (2021)

[25] Brombacher, E., Hackenberg, M., Kreutz, C., Binder, H., Treppner, M.: The performance of deep generative models for learning joint embeddings of single-cell multi-omics data. Front Mol Biosci 9, 962644 (2022)

[26] Hao, Y., Hao, S., Andersen-Nissen, E., Mauck, W.M., Zheng, S., Butler Lee, M.J., Wilk, A.J., Darby, C., Zager, M., Hoffman, P., Stoeckius, M., Papalexi, E., Mimitou, E.P., Jain, J., Srivastava, A., Stuart, T., Fleming, L.M., Yeung, B., Rogers, A.J., McElrath, J.M., Blish, C.A., Gottardo, R., Smibert, P., Satija, R.: Integrated analysis of multimodal single-cell data. Cell 184(13), 3573–3587 (2021)

[27] [27] Mourragui, S.M.C., Loog, M., van Nee, M., van de Wiel, M.A., Reinders, M.J.T., Wessels, L.F.A.: Percolate: an exponential family jive model to design dna-based predictors of drug response. bioRxiv (2022) https://www.biorxiv.org/content/early/2022/11/07/2022.09.11.507473.full.pdf. https://doi.org/10.1101/2022.09.11.507473

[28] [28] Eltager, M., Abdelaal, T., Charrout, M., Mahfouz, A., Reinders, M.J.T., Makrodimitris, S.: Benchmarking variational autoencoders on cancer transcriptomics data. bioRxiv (2023) https://www.biorxiv.org/content/early/2023/02/10/2023.02.09.527832.full.pdf. https://doi.org/10.1101/2023.02.09.527832

[29] Kingma, D.P., Rezende, D.J., Mohamed, S., Welling, M.: Semi-Supervised Learning with Deep Generative Models. arXiv (2014). https://doi.org/10.48550/ARXIV.1406.5298. https://arxiv.org/abs/1406.5298

[30] Gille, C., Guyard, F., Barlaud, M.: Semi-supervised classification using a supervised autoencoder for biomedical applications. arXiv (2022). https://doi.org/10.48550/ARXIV.2208.10315. https://arxiv.org/abs/2208.10315

[31] Breiman, L.: Heuristics of instability and stabilization in model selection. The Annals of Statistics 24(6), 2350–2383 (1996). Accessed 2023-03-14

[32] Žitnik, M., Zupan, B.: Data fusion by matrix factorization. IEEE Transactions on Pattern Analysis and Machine Intelligence 37(1), 41–53 (2015). https://doi.org/10.1109/TPAMI.2014.2343973

[33] Marcellino, L., Navarra, G.: A gpu-accelerated svd algorithm, based on qr factorization and givens rotations, for dwi denoising. In: 2016 12th International Conference on Signal-Image Technology & Internet-Based Systems (SITIS), pp. 699–704 (2016). https://doi.org/10.1109/SITIS.2016.117

[34] Tucker, G., Lawson, D., Gu, S., Maddison, C.J.: Doubly Reparameterized Gradient Estimators for Monte Carlo Objectives (2018)

[35] Chang, K., Creighton, C.J., Davis, C., Donehower, L., Drummond, J., Wheeler, D., et al.: The cancer genome atlas pan-cancer analysis project. Nature Genetics 45(10), 1113–1120 (2013). https://doi.org/10.1038/ng.2764

[36] The Cancer Genome Atlas: Pan-Cancer Atlas dataset: gene expression RNAseq - Batch effects normalized mRNA data. The Cancer Genome Atlas. Accessed: 20-04-2021 (2016)

[37] The Cancer Genome Atlas: Pan-Cancer Atlas dataset: copy number (gene-level) - gene-level copy number (gistic2). The Cancer Genome Atlas. Accessed: 20-04-2021 (2016)

[38] Mermel, C.H., Schumacher, S.E., Hill, B., Meyerson, M.L., Beroukhim, R., Getz, G.: Gistic2.0 facilitates sensitive and confident localization of the targets of focal somatic copy-number alteration in human cancers. Genome Biology 12(4) (2011). https://doi.org/10.1186/gb-2011-12-4-r41

[39] The Cancer Genome Atlas: Pan-Cancer Atlas dataset: DNA methylation - DNA methylation (Methylation450K). The Cancer Genome Atlas. Accessed: 20-04-2021 (2016)

[40] The Cancer Genome Atlas: Pan-Cancer Atlas dataset: Phenotype - Curated clinical data. The Cancer Genome Atlas. Accessed: 08-06-2021 (2018)

[41] Liu, J., Lichtenberg, T., Hoadley, K.A., Poisson, L.M., Lazar, A.J., Cherniack, A.D., Kovatich, A.J., Benz, C.C., Levine, D.A., Lee, A.V., Omberg, L., Wolf, D.M., Shriver, C.D., Thorsson, V., Hu, H., Caesar-Johnson, S.J., Demchok, J.A., Felau, I., Kasapi, M., Ferguson, M.L., Hutter, C.M., Sofia, H.J., Tarnuzzer, R., Wang, Z., Yang, L., Zenklusen, J.C., Zhang, J.J., Chudamani, S., Liu, J., Lolla, L., Naresh, R., Pihl, T., Sun, Q., Wan, Y., Wu, Y., Cho, J., DeFreitas, T., Frazer, S., Gehlenborg, N., Getz, G., Heiman, D.I., Kim, J., Lawrence, M.S., Lin, P., Meier, S., Noble, M.S., Saksena, G., Voet, D., Zhang, H., Bernard, B., Chambwe, N., Dhankani, V., Knijnenburg, T., Kramer, R., Leinonen, K., Liu, Y., Miller, M., Reynolds, S., Shmulevich, I., Thorsson, V., Zhang, W., Akbani, R., Broom, B.M., Hegde, A.M., Ju, Z., Kanchi, R.S., Korkut, A., Li, J., Liang, H., Ling, S., Liu, W., Lu, Y., Mills, G.B., Ng, K.S., Rao, A., Ryan, M., Wang, J., Weinstein, J.N., Zhang, J., Abeshouse, A., Armenia, J., Chakravarty, D., Chatila, W.K., de Bruijn, I., Gao, J., Gross, B.E., Heins, Z.J., Kundra, R., La, K., Ladanyi, M., Luna, A., Nissan, M.G., Ochoa, A., Phillips, S.M., Reznik, E., Sanchez-Vega, F., Sander, C., Schultz, N., Sheridan, R., Sumer, S.O., Sun, Y., Taylor, B.S., Wang, J., Zhang, H., Anur, P., Peto, M., Spellman, P., Benz, C., Stuart, J.M., Wong, C.K., Yau, C., Hayes, D.N., Parker, J.S., Wilkerson, M.D., Ally, A., Balasundaram, M., Bowlby, R., Brooks, D., Carlsen, R., Chuah, E., Dhalla, N., Holt, R., Jones, S.J.M., Kasaian, K., Lee, D., Ma, Y., Marra, M.A., Mayo, M., Moore, R.A., Mungall, A.J., Mungall, K., Robertson, A.G., Sadeghi, S., Schein, J.E., Sipahimalani, P., Tam, A., Thiessen, N., Tse, K., Wong, T., Berger, A.C., Beroukhim, R., Cherniack, A.D., Cibulskis, C., Gabriel, S.B., Gao, G.F., Ha, G., Meyerson, M., Schumacher, S.E., Shih, J., Kucherlapati, M.H., Kucherlapati, R.S., Baylin, S., Cope, L., Danilova, L., Bootwalla, M.S., Lai, P.H., Maglinte, D.T., Van Den Berg, D.J., Weisenberger, D.J., Auman, J.T., Balu, S., Bodenheimer, T., Fan, C., Hoadley, K.A., Hoyle, A.P., Jefferys, S.R., Jones, C.D., Meng, S., Mieczkowski, P.A., Mose, L.E., Perou, A.H., Perou, C.M., Roach, J., Shi, Y., Simons, J.V., Skelly, T., Soloway, M.G., Tan, D., Veluvolu, U., Fan, H., Hinoue, T., Laird, P.W., Shen, H., Zhou, W., Bellair, M., Chang, K., Covington, K., Creighton, C.J., Dinh, H., Doddapaneni, H., Donehower, L.A., Drummond, J., Gibbs, R.A., Glenn, R., Hale, W., Han, Y., Hu, J., Korchina, V., Lee, S., Lewis, L., Li, W., Liu, X., Morgan, M., Morton, D., Muzny, D., Santibanez, J., Sheth, M., Shinbro, E., Wang, L., Wang, M., Wheeler, D.A., Xi, L., Zhao, F., Hess, J., Appelbaum, E.L., Bailey, M., Cordes, M.G., Ding, L., Fronick, C.C., Fulton, L.A., Fulton, R.S., Kandoth, C., Mardis, E.R., McLellan, M.D., Miller, C.A., Schmidt, H.K., Wilson, R.K., Crain, D., Curley, E., Gardner, J., Lau, K., Mallery, D., Morris, S., Paulauskis, J., Penny, R., Shelton, C., Shelton, T., Sherman, M., Thompson, E., Yena, P., Bowen, J., Gastier-Foster, J.M., Gerken, M., Leraas, K.M., Lichtenberg, T.M., Ramirez, N.C., Wise, L., Zmuda, E., Corcoran, N., Costello, T., Hovens, C., Carvalho, A.L., de Carvalho, A.C., Fregnani, J.H., Longatto-Filho, A., Reis, R.M., Scapulatempo-Neto, C., Silveira, H.C.S., Vidal, D.O., Burnette, A., Eschbacher, J., Hermes, B., Noss, A., Singh, R., Anderson, M.L., Castro, P.D., Ittmann, M., Huntsman, D., Kohl, B., Le, X., Thorp, R., Andry, C., Duffy, E.R., Lyadov, V., Paklina, O., Setdikova, G., Shabunin, A., Tavobilov, M., McPherson, C., Warnick, R., Berkowitz, R., Cramer, D., Feltmate, C., Horowitz, N., Kibel, A., Muto, M., Raut, C.P., Malykh, A., Barnholtz-Sloan, J.S., Barrett, W., Devine, K., Fulop, J., Ostrom, Q.T., Shimmel, K., Wolinsky, Y., Sloan, A.E., De Rose, A., Giuliante, F., Goodman, M., Karlan, B.Y., Hagedorn, C.H., Eckman, J., Harr, J., Myers, J., Tucker, K., Zach, L.A., Deyarmin, B., Hu, H., Kvecher, L., Larson, C., Mural, R.J., Somiari, S., Vicha, A., Zelinka, T., Bennett, J., Iacocca, M., Rabeno, B., Swanson, P., Latour, M., Lacombe, L., tu, B., Bergeron, A., McGraw, M., Staugaitis, S.M., Chabot, J., Hibshoosh, H., Sepulveda, A., Su, T., Wang, T., Potapova, O., Voronina, O., Desjardins, L., Mariani, O., Roman-Roman, S., Sastre, X., Stern, M.H., Cheng, F., Signoretti, S., Berchuck, A., Bigner, D., Lipp, E., Marks, J., McCall, S., McLendon, R., Secord, A., Sharp, A., Behera, M., Brat, D.J., Chen, A., Delman, K., Force, S., Khuri, F., Magliocca, K., Maithel, S., Olson, J.J., Owonikoko, T., Pickens, A., Ramalingam, S., Shin, D.M., Sica, G., Van Meir, E.G., Zhang, H., Eijckenboom, W., Gillis, A., Korpershoek, E., Looijenga, L., Oosterhuis, W., Stoop, H., van Kessel, K.E., Zwarthoff, E.C., Calatozzolo, C., Cuppini, L., Cuzzubbo, S., DiMeco, F., Finocchiaro, G., Mattei, L., Perin, A., Pollo, B., Chen, C., Houck, J., Lohavanichbutr, P., Hartmann, A., Stoehr, C., Stoehr, R., Taubert, H., Wach, S., Wullich, B., Kycler, W., Murawa, D., Wiznerowicz, M., Chung, K., Edenfield, W.J., Martin, J., Baudin, E., Bubley, G., Bueno, R., De Rienzo, A., Richards, W.G., Kalkanis, S., Mikkelsen, T., Noushmehr, H., Scarpace, L., Girard, N., Aymerich, M., Campo, E.e E., Guillermo, A.L., Van Bang, N., Hanh, P.T., Phu, B.D., Tang, Y., Colman, H., Evason, K., Dottino, P.R., Martignetti, J.A., Gabra, H., Juhl, H., Akeredolu, T., Stepa, S., Hoon, D., Ahn, K., Kang, K.J., Beuschlein, F., Breggia, A., Birrer, M., Bell, D., Borad, M., Bryce, A.H., Castle, E., Chandan, V., Cheville, J., Copland, J.A., Farnell, M., Flotte, T., Giama, N., Ho, T., Kendrick, M., Kocher, J.P., Kopp, K., Moser, C., Nagorney, D., O’Brien, D., O’Neill, B.P., Patel, T., Petersen, G., Que, F., Rivera, M., Roberts, L., Smallridge, R., Smyrk, T., Stanton, M., Thompson, R.H., Torbenson, M., Yang, J.D., Zhang, L., Brimo, F., Ajani, J.A., Angulo Gonzalez, A.M., Behrens, C., Bondaruk, J., Broaddus, R., Czerniak, B., Esmaeli, B., Fujimoto, J., Gershenwald, J., Guo, C., Lazar, A.J., Logothetis, C., Meric-Bernstam, F., Moran, C., Ramondetta, L., Rice, D., Sood, A., Tamboli, P., Thompson, T., Troncoso, P., Tsao, A., Wistuba, I., Carter, C., Haydu, L., Hersey, P., Jakrot, V., Kaka-vand, H., Kefford, R., Lee, K., Long, G., Mann, G., Quinn, M., Saw, R., Scolyer, R., Shannon, K., Spillane, A., Stretch, J., Synott, M., Thompson, J., Wilmott, J., Al-Ahmadie, H., Chan, T.A., Ghossein, R., Gopalan, A., Levine, D.A., Reuter, V., Singer, S., Singh, B., Tien, N.V., Broudy, T., Mirsaidi, C., Nair, P., Drwiega, P., Miller, J., Smith, J., Zaren, H., Park, J.W., Hung, N.P., Kebebew, E., Linehan, W.M., Metwalli, A.R., Pacak, K., Pinto, P.A., Schiffman, M., Schmidt, L.S., Vocke, C.D., Wentzensen, N., Worrell, R., Yang, H., Moncrieff, M., Goparaju, C., Melamed, J., Pass, H., Botnariuc, N., Caraman, I., Cernat, M., Chemencedji, I., Clipca, A., Doruc, S., Gorincioi, G., Mura, S., Pirtac, M., Stancul, I., Tcaciuc, D., Albert, M., Alexopoulou, I., Arnaout, A., Bartlett, J., Engel, J., Gilbert, S., Parfitt, J., Sekhon, H., Thomas, G., Rassl, D.M., Rintoul, R.C., Bifulco, C., Tamakawa, R., Urba, W., Hayward, N., Timmers, H., Antenucci, A., Facciolo, F., Grazi, G., Marino, M., Merola, R., de Krijger, R., Gimenez-Roqueplo, A.P.e A., Chevalier, S., McKercher, G., Birsoy, K., Barnett, G., Brewer, C., Farver, C., Naska, T., Pennell, N.A., Raymond, D., Schilero, C., Smolenski, K., Williams, F., Morrison, C., Borgia, J.A., Liptay, M.J., Pool, M., Seder, C.W., Junker, K., Omberg, L., Dinkin, M., Manikhas, G., Alvaro, D., Bragazzi, M.C., Cardinale, V., Carpino, G., Gaudio, E., Chesla, D., Cottingham, S., Dubina, M., Moiseenko, F., Dhanasekaran, R., Becker, K.F., Janssen, K.P., Slotta-Huspenina, J., Abdel-Rahman, M.H., Aziz, D., Bell, S., Cebulla, C.M., Davis, A., Duell, R., Elder, J.B., Hilty, J., Kumar, B., Lang, J., Lehman, N.L., Mandt, R., Nguyen, P., Pilarski, R., Rai, K., Schoenfield, L., Senecal, K., Wakely, P., Hansen, P., Lechan, R., Powers, J., Tischler, A., Grizzle, W.E., Sexton, K.C., Kastl, A., Henderson, J., Porten, S., Waldmann, J., Fassnacht, M., Asa, S.L., Schadendorf, D., Couce, M., Graefen, M., Huland, H., Sauter, G., Schlomm, T., Simon, R., Tennstedt, P., Olabode, O., Nelson, M., Bathe, O., Carroll, P.R., Chan, J.M., Disaia, P., Glenn, P., Kelley, R.K., Landen, C.N., Phillips, J., Prados, M., Simko, J., Smith-McCune, K., VandenBerg, S., Roggin, K., Fehrenbach, A., Kendler, A., Sifri, S., Steele, R., Jimeno, A., Carey, F., Forgie, I., Mannelli, M., Carney, M., Hernandez, B., Campos, B., Herold-Mende, C., Jungk, C., Unterberg, A., von Deimling, A., Bossler, A., Galbraith, J., Jacobus, L., Knudson, M., Knutson, T., Ma, D., Milhem, M., Sigmund, R., Godwin, A.K., Madan, R., Rosenthal, H.G., Adebamowo, C., Adebamowo, S.N., Boussioutas, A., Beer, D., Giordano, T., Mes-Masson, A.M., Saad, F., Bocklage, T., Landrum, L., Mannel, R., Moore, K., Moxley, K., Postier, R., Walker, J., Zuna, R., Feldman, M., Valdivieso, F., Dhir, R., Luketich, J., Mora Pinero, E.M., Quintero-Aguilo, M., Carlotti, C.G., Dos Santos, J.S., Kemp, R., Sankarankuty, A., Tirapelli, D., Catto, J., Agnew, K., Swisher, E., Creaney, J., Robinson, B., Shelley, C.S., Godwin, E.M., Kendall, S., Shipman, C., Bradford, C., Carey, T., Haddad, A., Moyer, J., Peterson, L., Prince, M., Rozek, L., Wolf, G., Bowman, R., Fong, K.M., Yang, I., Korst, R., Rathmell, W.K., Fantacone-Campbell, J.L., Hooke, J.A., Kovatich, A.J., Shriver, C.D., DiPersio, J., Drake, B., Govindan, R., Heath, S., Ley, T., Van Tine, B., Westervelt, P., Rubin, M.A., Lee, J.I., Aredes, N.D., Mariamidze, A.: An Integrated TCGA Pan-Cancer Clinical Data Resource to Drive High-Quality Survival Outcome Analytics. Cell 173(2), 400–416 (2018)

[42] Akaike, H.: In: Parzen, E., Tanabe, K., Kitagawa, G. (eds.) Information Theory and an Extension of the Maximum Likelihood Principle, pp. 199–213. Springer, New York, NY (1998). https://doi.org/10.1007/978-1-4612-1694-015. https://doi.org/10.1007/978-1-4612-1694-015

[43] Stone, M.: An asymptotic equivalence of choice of model by cross-validation and akaike’s criterion. Journal of the Royal Statistical Society. Series B (Methodological) 39(1), 44–47 (1977). Accessed 2023-02-01

