## Supplementary Tables for "An in-depth comparison of linear and non-linear joint embedding methods for bulk and single-cell multi-omics"

Table S1: Mean imputation log-likelihood of gene expression (GE) from DNA methylation (ME) and vice-versa on the TCGA dataset. Higher log-likelihood is better irrespective of the sign. The values for both the validation and the test data are listed.

| | VALIDATION SET | | TEST SET | |
|---|---|---|---|---|
| model | GE from ME | ME from GE | GE from ME | ME from GE |
| GLM | -4,193.94 | 1,950.64 | -4,201.91 | 1,805.08 |
| MCIA | -5,000.33 | 4,323.83 | -4,980.13 | 4,335.12 |
| MOFA+ | -5,376.41 | 4,340.35 | -5,368.54 | 4,306.57 |
| CGVAE | -4,162.21 | 5,119.92 | -4,149.46 | 5,062.66 |
| ccVAE | -6,573.50 | 2,183.08 | -6,682.89 | 2,220.33 |
| PoE | -3,763.50 | 5,517.53 | -3,787.29 | 5,493.66 |
| MoE | -3,861.87 | 5,183.17 | -3,787.29 | 5183.18 |

Table S2: Mean imputation log-likelihood of gene expression (GE) from copy number (CNV) and vice-versa on the TCGA dataset. Higher log-likelihood is better irrespective of the sign. The values for both the validation and the test data are listed.

| | VALIDATION SET | | TEST SET | |
|---|---|---|---|---|
| model | GE from CNV | CNV from GE | GE from CNV | CNV from GE |
| GLM | -4,193.94 | 1,950.64 | -4,201.91 | 1,805.08 |
| MCIA | -11,571.34 | 4,323.83 | -9,880.65 | 4,335.12 |
| MOFA+ | -5,376.41 | 4,340.35 | -5,368.54 | 4,306.57 |
| CGVAE | -4,162.21 | 5,119.92 | -4,149.46 | 5,062.66 |
| ccVAE | -6,573.50 | 2,183.08 | -6,682.89 | 2,220.33 |
| PoE | -3,763.50 | 5,517.53 | -3,787.29 | 5,493.66 |
| MoE | -3,861.87 | 5,183.17 | -3,787.29 | 5183.18 |

Table S3: Optimal hyperparamaters for TCGA dataset based on validation loss

| dataset | model | latent dimension | encoder layers | learning rate | dropout | batch normalization | K |
|---|---|---|---|---|---|---|---|
| GE + ME | MCIA | 20 | - | - | - | - | - |
| | MOFA+ | 47 | - | - | - | - | - |
| | CGVAE | 64 | 256-256 | 0.001 | 0% | Yes | - |
| | ccVAE | 64 | 256-256 | 0.001 | 0% | Yes | - |
| | PoE | 64 | 256 | 0.0001 | 10% | No | - |
| | MoE | 32 | 256-256 | 0.0001 | 0% | No | 10 |
| GE + CNV | MCIA | 16 | - | - | - | - | - |
| | MOFA+ | 60 | - | - | - | - | - |
| | CGVAE | 64 | 256-128 | 0.0001 | 0% | No | - |
| | ccVAE | 64 | 256 | 0.001 | 0% | Yes | - |
| | PoE | 64 | 256 | 0.001 | 10% | Yes | - |
| | MoE | 32 | 256-128 | 0.0001 | 10% | No | 20 |

Table S4: Validation and test performance (Matthews Correlation Coefficient - MCC) of neural networks that predict tumor type from an input omic profile on the TCGA and (level 2) cell type from an input single-cell profile on the CITE-Seq data. MCC of 0 corresponds to random guessing and MCC of 1 to perfect classification.

| Dataset | Data type | validation MCC | test MCC |
|---|---|---|---|
| TCGA | GE | 0.968 | 0.958 |
| TCGA | ME | 0.962 | 0.955 |
| TCGA | CNV | 0.406 | 0.440 |
| CITE-Seq | RNA | 0.929 | 0.915 |
| CITE-Seq | ADT | 0.936 | 0.904 |

Table S5: Survival analysis GE+ME

| Model | Modality | AIC | #factors | significant factors | % significant factors |
|---|---|---|---|---|---|
| covariates only | - | 24,316.76 | - | - | - |
| PCA | GE | 23,936.77 | 32 | 13 | 41% |
| | ME | 24,034.69 | 32 | 15 | 47% |
| | GE+ME | 23,858.65 | 64 | 31 | 48% |
| MCIA | GE | 24,015.08 | 20 | 10 | 50% |
| | ME | 24,024.65 | 20 | 10 | 50% |
| | GE+ME | 24,020.04 | 40 | 6 | 15% |
| MOFA+ | GE | 23,904.53 | 47 | 13 | 28% |
| | ME | 23,937.95 | 47 | 15 | 32% |
| | GE+ME | 23,918.48 | 94 | 10 | 11% |
| CGVAE | GE | 23,890.94 | 64 | 15 | 23% |
| | ME | 23,883.19 | 64 | 15 | 23% |
| | GE+ME | 23,870.96 | 128 | 20 | 16% |
| ccVAE | GE+ME | 23,860.87 | 64 | 17 | 27% |
| PoE | GE+ME | 23,906.31 | 64 | 21 | 33% |
| MoE | GE | 23,966.63 | 32 | 14 | 44% |
| | ME | 23,938.41 | 32 | 6 | 19% |
| | GE+ME | 23,911.54 | 64 | 11 | 18% |

Table S6: Survival analysis GE+CNV

| Model | Modality | AIC | #factors | significant factors | % significant factors |
|---|---|---|---|---|---|
| covariates only | - | 24,316.76 | - | - | - |
| PCA | GE | 23,936.77 | 32 | 13 | 41% |
| | CNV | 24,283.20 | 32 | 9 | 28% |
| | GE+CNV | 23,947.96 | 64 | 17 | 27% |
| MCIA | GE | 24,211.88 | 16 | 6 | 38% |
| | CNV | 24,355.63 | 16 | 0 | 0% |
| | GE+CNV | 24,306.60 | 32 | 6 | 19% |
| MOFA+ | GE | 23,992.16 | 60 | 21 | 35% |
| | CNV | 24,443.58 | 60 | 0 | 0% |
| | GE+CNV | 24,297.19 | 120 | 17 | 14% |
| CGVAE | GE | 24,414.22 | 64 | 0 | 0% |
| | CNV | 24,253.06 | 64 | 3 | 5% |
| | GE+CNV | 24,525.55 | 128 | 0 | 0% |
| ccVAE | GE+CNV | 23,991.02 | 64 | 16 | 25% |
| PoE | GE+CNV | 23,920.78 | 64 | 19 | 30% |
| MoE | GE | 24,237.80 | 32 | 12 | 38% |
| | CNV | 24,262.78 | 32 | 3 | 9% |
| | GE+CNV | 24,204.19 | 64 | 14 | 22% |

Table S7: Optimal hyperparamaters for CITE-Seq dataset based on validation loss

| model | latent dimension | encoder layers | learning rate | dropout | batch normalization | K |
|---|---|---|---|---|---|---|
| MCIA | - | - | - | - | - | - |
| MOFA+ | 63 | - | - | - | - | - |
| CGVAE | 64 | 128 | 0.0001 | 0% | No | - |
| ccVAE | 64 | 256 | 0.0001 | 0% | No | - |
| PoE | 64 | - | 0.001 | 10% | No | - |
| MoE | 32 | 256-128 | 0.0001 | 0% | No | 10 |

Table S8: Mean imputation log-likelihood of gene expression (RNA) from protein expression (ADT) and vice-versa on the PBMC dataset. Higher log-likelihood is better irrespective of the sign. The values for both the validation and the test data are listed.

| | VALIDATION SET | | TEST SET | |
|---|---|---|---|---|
| model | RNA from ADT | ADT from RNA | RNA from ADT | ADT from RNA |
| GLM | -913.67 | -232.45 | -968.67 | -225.24 |
| MOFA+ | -903.02 | -253.74 | -961.29 | -267.406 |
| CGVAE | -884.43 | -235.855 | -940.139 | -227.70 |
| ccVAE | -904.33 | -136357.16 | -957.90 | -186,784.09 |
| PoE | -879.96 | -244.32 | -984.86 | -518.98 |
| MoE | -865.20 | -225.56 | -918.26 | -220.64 |

Table S9: Cell type classification performance (Matthews Correlation Coefficient) on the CITE-Seq dataset

| Model | Modality | SVM MCC | MLP MCC |
|---|---|---|---|
| PCA | RNA | 0.759 | 0.808 |
| | ADT | 0.722 | 0.775 |
| | RNA+ADT | 0.818 | 0.871 |
| MOFA+ | RNA | 0.764 | 0.872 |
| | ADT | 0.751 | 0.833 |
| | RNA+ADT | 0.797 | 0.889 |
| CGVAE | RNA | 0.612 | 0.748 |
| | ADT | 0.581 | 0.713 |
| | RNA+ADT | 0.633 | 0.805 |
| ccVAE | RNA | 0.579 | 0.750 |
| | ADT | 0.661 | 0.735 |
| | RNA+ADT | 0.666 | 0.744 |
| PoE | RNA | 0.835 | 0.874 |
| | ADT | 0.820 | 0.828 |
| | RNA+ADT | 0.830 | 0.840 |
| MoE | RNA | 0.749 | 0.826 |
| | ADT | 0.757 | 0.807 |
| | RNA+ADT | 0.826 | 0.870 |

Table S10: Percent agreement of the MLP predictions using a measured profile and an imputed profile using a single-modal or multi-modal classifier trained solely on real data.

| | MOFA+ | PoE | MoE |
|---|---|---|---|
| RNA from ADT (unimodal) | 6.2% | 71.3% | 77.5% |
| ADT from RNA (unimodal) | 77.3% | 74.7% | 73.3% |
| RNA from ADT (multimodal) | 11.5% | 83.6% | 86.2% |
| ADT from RNA (multimodal) | 89.1% | 89.6% | 88.7% |
